# The mechanism of Atg15-mediated membrane disruption in autophagy

**DOI:** 10.1101/2023.06.26.546516

**Authors:** Yoko Kagohashi, Michiko Sasaki, Alexander I May, Tomoko Kawamata, Yoshinori Ohsumi

## Abstract

Autophagy is a lysosomal/vacuolar delivery system that isolates and degrades cytoplasmic material. Following delivery by autophagosomes, cytoplasmic components are released into the vacuole within an autophagic body (AB), which is a single-membrane structure derived from the inner membrane of the autophagosome. This membrane must be disrupted for degradation of the cytoplasmic cargo to occur. The vacuolar proteases Pep4 and Prb1, as well as the lipase Atg15, are known to be necessary for this process, but the mechanistic underpinnings remain unclear. In this study, we establish a system to detect lipase activity in the vacuole and use it to show that Atg15 is the sole vacuolar phospholipase and that Pep4 and Prb1 are required for the activation of Atg15 lipase function, which occurs following delivery of Atg15 to the vacuole by the MVB pathway. In vitro experiments also reveal that Atg15 is a B-type phospholipase of broad substrate specificity that is likely implicated in the disruption of a range of membranes delivered to the vacuole. Further, we use isolated ABs to demonstrate that Atg15 alone is able to disrupt AB membranes.

## Introduction

Intracellular homeostasis is maintained by continuous synthesis and degradation of cellular components. Autophagy, a major intracellular degradation system highly conserved among eukaryotes, plays a key role in multiple metabolic and cellular processes (Ohsumi, 2014; Parzych and Klionsky, 2014). Disruption of this degradation results in the accumulation of cargo materials within the vacuole/lysosome and the inability to adapt to starvation conditions in yeast (Takeshige et al., 1992), or various lysosomal storage diseases in mammals (Parkinson-Lawrence et al., 2010). Yeast provides an excellent system to elucidate the molecular mechanisms of degradation in the vacuole because of its genetic tractability and ease of biochemical experimentation. We have previously shown that autophagy-dependent RNA degradation in the vacuole is mediated by a T2-type RNase, Rny1, and the nucleotidase Pho8 (Huang et al., 2015).

Upon induction of autophagy in yeast, a small membrane sack appears, which elongates and subsequently closes to form a double membrane vesicle known as an autophagosome. The autophagosome sequesters a portion of cytoplasm before its outer membrane fuses with the vacuole, releasing the inner membrane structure, called the autophagic body (AB), into the vacuolar lumen. ABs are immediately disintegrated, releasing their cargoes for degradation by vacuolar hydrolases into compounds that can be recycled for new synthesis. Therefore, the first critical step of degradation following delivery to the vacuole is the disruption of the AB membrane to expose cargoes to hydrolytic vacuolar enzymes.

Vacuolar proteinase A (Pep4) and proteinase B (Prb1) have been shown to be necessary for AB disruption in yeast (Takeshige et al., 1992). Pep4 is an aspartic endoprotease that triggers the protease cascade, resulting in the activation of many hydrolases that amplify the degradative capacity of the vacuole (Ammerer et al., 1986). Prb1 is a serine endoprotease that is activated by Pep4 (Mechler et al., 1988; Moehle et al., 1989). These two proteases are involved in the activation of other hydrolases including carboxypeptidase Y (CPY), aminopeptidase I (Ape1), and Pho8 (Hecht et al., 2014; Parzych and Klionsky, 2014). Treatment with a serine protease inhibitor, phenylmethylsulfonyl fluoride (PMSF), also inhibits AB disruption (Takeshige et al., 1992). It is not clear why these vacuolar proteases are essential for the degradation of the limiting membrane of ABs. Given that electron microscopy indicates that AB membranes contain few proteins (Baba et al., 1995), it is unlikely that vacuolar proteases directly disrupt AB membranes.

Degradation of glycerophospholipids, which are the main components of lipid bilayers, may be essential for membrane disruption. In yeast, *ATG15* (identified initially as *AUT5*/*CVT1*) encodes a vacuolar phospholipase with a single transmembrane domain (Epple et al., 2001; Teter et al., 2001). During synthesis, Atg15 undergoes glycosylation and is then transported to the vacuole via the multi-vesicular body (MVB) pathway (Epple et al., 2001). The lack of Atg15 causes accumulation of ABs in the vacuole (Epple et al., 2001). Atg15 is also involved in the disruption of other transport vesicles such as those of the cytoplasm-to-vacuole targeting (Cvt) pathway, internal vesicles of MVB pathway, and microautophagic bodies (Epple et al., 2003; Oku et al., 2017; Teter et al., 2001). However, the active form of Atg15 is yet to be characterized.

Phospholipases are classified into four groups based on the specific cleavage of ester bonds: phospholipase A cleaves the acyl ester bond at either the *sn-1* (phospholipase A1) or *sn-2* (phospholipase A2) position of a phospholipid, liberating free fatty acid (FFA) and lysophospholipid (LPL); phospholipase B cleaves acyl chains at the *sn-1* and *sn-2* positions of phospholipid, releasing two free FFAs; phospholipase C cleaves the glycerophosphate bond; and phospholipase D removes the polar head group (Stahelin, 2016).

Ramya & Rajasekharan have reported that purified Atg15 from microsomal membrane fractions cleaves the ester bond at the *sn-1* position of phospholipids and has high activity toward phosphatidylserine (PS), moderate activity toward phosphatidylethanolamine (PE) and cardiolipin (CL), but almost no activity toward phosphatidic acid (PA), phosphatidylinositol (PI), phosphatidylcholine (PC), phosphatidylglycerol (PG), or lysophospholipid (LPL) (Ramya and Rajasekharan, 2016). On the other hand, van Zutphen et al. showed that Atg15 acts as a lipase that processes the triglyceride (TG) class of neutral lipids (van Zutphen et al., 2014). It remains unclear whether these phospholipase A1 activities and reported substrate specificities are sufficient to explain the process of AB disruption. In this study, we assess Atg15 and Pep4/Prb1 activities to clarify the long-standing question of how AB membranes are disrupted.

## Results

### *ATG15* confers vacuolar lipase activity

*ATG15* encodes a protein with a single transmembrane domain at the N-terminus (Fig 1A). The residues 330-334 constitute a GXSXG lipase consensus motif, and substitution of the serine 332 with alanine (S332A) results in a defect in AB disruption (Epple et al., 2001). Since there exists several lipases in various compartments (Ramya and Rajasekharan, 2016), we first set out to develop an in vitro assay system that can detect lipase activity strictly within the vacuole. Vacuoles were purified from wild-type (WT) cells as previously reported (Ohsumi and Anraku, 1981) and disrupted by freeze-thaw cycles to yield a vacuolar lysate (Fig 1B, Fig. EV1A). To monitor lipase activity, we used NBD-PE, which is PE labeled with nitrobenzoxadiazole (NBD) on the methyl end of the FFA moiety at *sn-2* (Fig 1B). Lipase mediated hydrolysis of the caboxylic ester of NBD-PE at the *sn-1* position (Ramya and Rajasekharan, 2016) would produce only NBD-labeled lysophosphatidylethanolamine (NBD-LPE). However, incubation of NBD-PE with vacuolar lysate from wild type cells yielded both NBD-LPE and NBD-labeled free FFAs (NBD-FFA; Fig 1C lane 2). These represent hydrolysis of the NBD-PE acyl ester linkage at *sn-1* (NBD-LPE) or the *sn-2* ester bond of NBD-PE (NBD-FFA); alternatively, further hydrolysis of NBD-LPE can also yield NBD-FFA. These results clearly indicate that vacuolar lysate is able to release FFAs by hydrolysis of *sn-1* and *sn-2*, suggesting phospholipase A1 and A2, or B-type activity.

**Figure 1.**
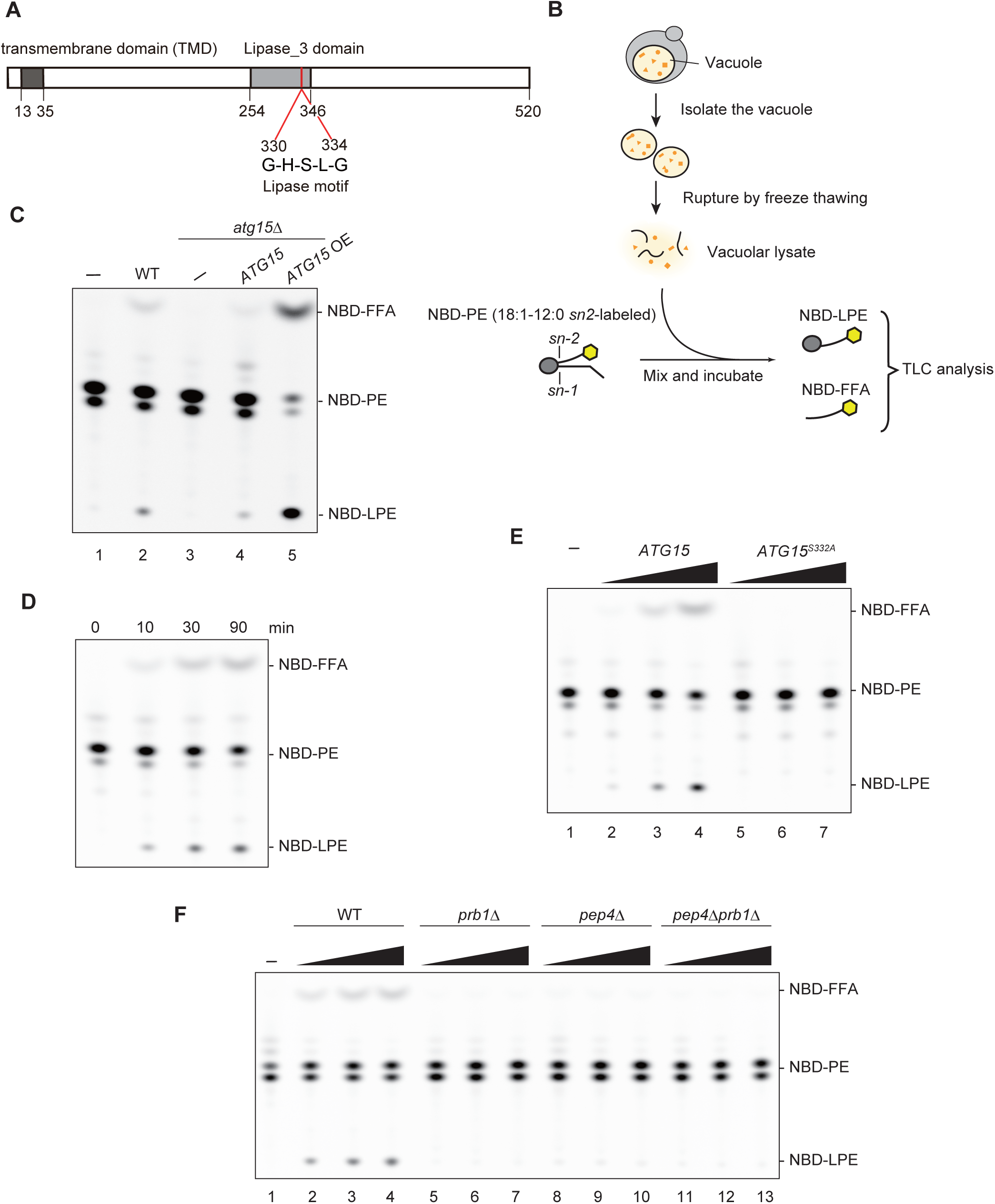
Vacuolar lipase activity derived from Atg15 depends on Pep4/Prb1. (A) Schematic diagram of Atg15. The transmembrane domain (TMD) and a lipase consensus motif are shown. (B) Schematic overview of the in vitro assay system used to measure vacuolar lipase activity. (C) Lipase activities of vacuolar lysates from WT, *atg1*5Δ, and *atg15*Δ cells harboring *ATG15* from either single or multi-copy (OE) plasmids. The amount of lysate used for assay was normalized by the amount of the vacuolar membrane protein, Pho8 (See Fig EV1B). (D) Lipase activity of vacuolar lysates from Atg15OE cells. (E) Lipase activities of vacuolar lysate from Atg15OE cells. Each vacuolar lysate was added at a volume ratio of 1:5:25. The amount of the lysates used for this assay was shown in Fig EV1C. (F) Lipase activities of vacuolar lysates from WT, *prb1*Δ, *pep4*Δ, and *pep4*Δ *prb1*Δ cells. Each vacuolar lysate was added at a volume ratio of 1:2:4. The amount of the lysates used for this assay was shown in Fig EV1D.

Next, we investigated the relationship between this lipase activity and Atg15. Vacuolar lysate from *atg15*Δ cells generated neither NBD-FFA nor NBD-LPE (Fig 1C lane 3 and Fig EV1B). These products reappeared when NBD-PE was incubated in the presence of lysates from *atg15*Δ cells expressing *ATG15* from a plasmid (Fig 1C lanes 4 and 5 and Fig EV1B). Product formation was increased when *ATG15* expression was elevated using a multicopy plasmid. Further, hydrolysis progressed with incubation time (Fig 1D) and accelerated with the amount of the lysate added (Fig 1E lanes 2-4). Meanwhile, the lysates from cells expressing Atg15^S332A^ exhibited no hydrolytic activity at all (Fig 1E lanes 5-7 and Fig EV1C). These in vitro data indicate that Atg15 is the sole source of phospholipase activity in the vacuole.

### Atg15 is activated by vacuolar proteases

Previously, we reported that the deletion of the vacuolar proteases, Pep4 and Prb1, results in accumulation of ABs in the vacuole (Takeshige et al., 1992). We next asked how vacuolar lipase activity relates to these vacuolar proteases. Using vacuolar lysates from *prb1*Δ, *pep4*Δ, and *pep4*Δ*prb1*Δ cells, we performed the lipase assay and found that lipase activity was completely absent in cells lacking either Prb1 or Pep4 (Fig 1F lanes 5-13 and Fig EV1D). Given that Pep4 proteolytically activates Prb1 (Mechler et al., 1988; Moehle et al., 1989), Prb1 may act directly to activate Atg15, consistent with our previous study that disruption of ABs is inhibited by treatment with PMSF (Takeshige et al., 1992). However, it has been reported that Prb1 retains catalytic activity even in the absence of Pep4 activity (Mechler et al., 1988; Moehle et al., 1989). We therefore decided to employ a double deletion mutant of Pep4 and Prb1 to ensure complete ablation of endoprotease activity. Even when Atg15 was overexpressed in *pep4*Δ *prb1*Δ cells, no lipase activity was detected (Fig EV1E and F). It is therefore likely that these proteases are involved in the processing of Atg15 to an active form.

### The vacuolar luminal region of Atg15 is sufficient to produce lipase activity

Atg15 is synthesized in the endoplasmic reticulum (ER) and transported to the vacuole via the MVB pathway (Fig 2A). Residues 13-35 of Atg15 constitute a transmembrane domain, which acts as a signal sequence for the MVB pathway (Hirata et al., 2021). To detect Atg15, we expressed Atg15 C-terminally tagged with FLAG (Atg15-FLAG); after 1 h rapamycin treatment, vacuolar lysate samples were obtained and subjected to immunoblotting using α-FLAG antibodies. Atg15-FLAG expressed from a single copy plasmid was only faintly detected (Fig EV2A lane 3). We therefore decided to use cells expressing Atg15-FLAG from a multicopy plasmid (Fig EV2A lane 4). We confirmed that lipase activity is apparent in vacuolar lysates, with similar results to those shown in Fig 1C (Fig EV2B). In *pep4*Δ *prb1*Δ cells, a glycosylated full-length Atg15-FLAG band at 75 kDa, as well as heavily glycosylated smear bands, were observed, as previously reported (Fig 2B lane 2 and EV2C; Epple et al., 2001). In wild type cells, a single band slightly smaller than the 75 kDa band was also apparent, the amount of which was reduced due to processing at the C-terminus (Fig 2B lane 1 and EV2C). When lysates from *pep4*Δ *prb1*Δ cells were treated with endoglycosidase H, the 75 kDa band and smear band converged to one lower band (Fig 2B lane 4). Further, compared with this band, a single band of apparently smaller molecular mass appeared in wild type lysates (Fig 2B lane 3). These observed differences in mobility of Atg15-FLAG suggests that the N-terminus of all Atg15 molecules was cleaved by Pep4/Prb1 since all observed bands have an intact C-terminus. Atg15 cleavage by Pep4/Prb1 most likely occurs at an N-terminal region just after transmembrane domain immediately following delivery of the Atg15 protein to the vacuole.

**Figure 2.**
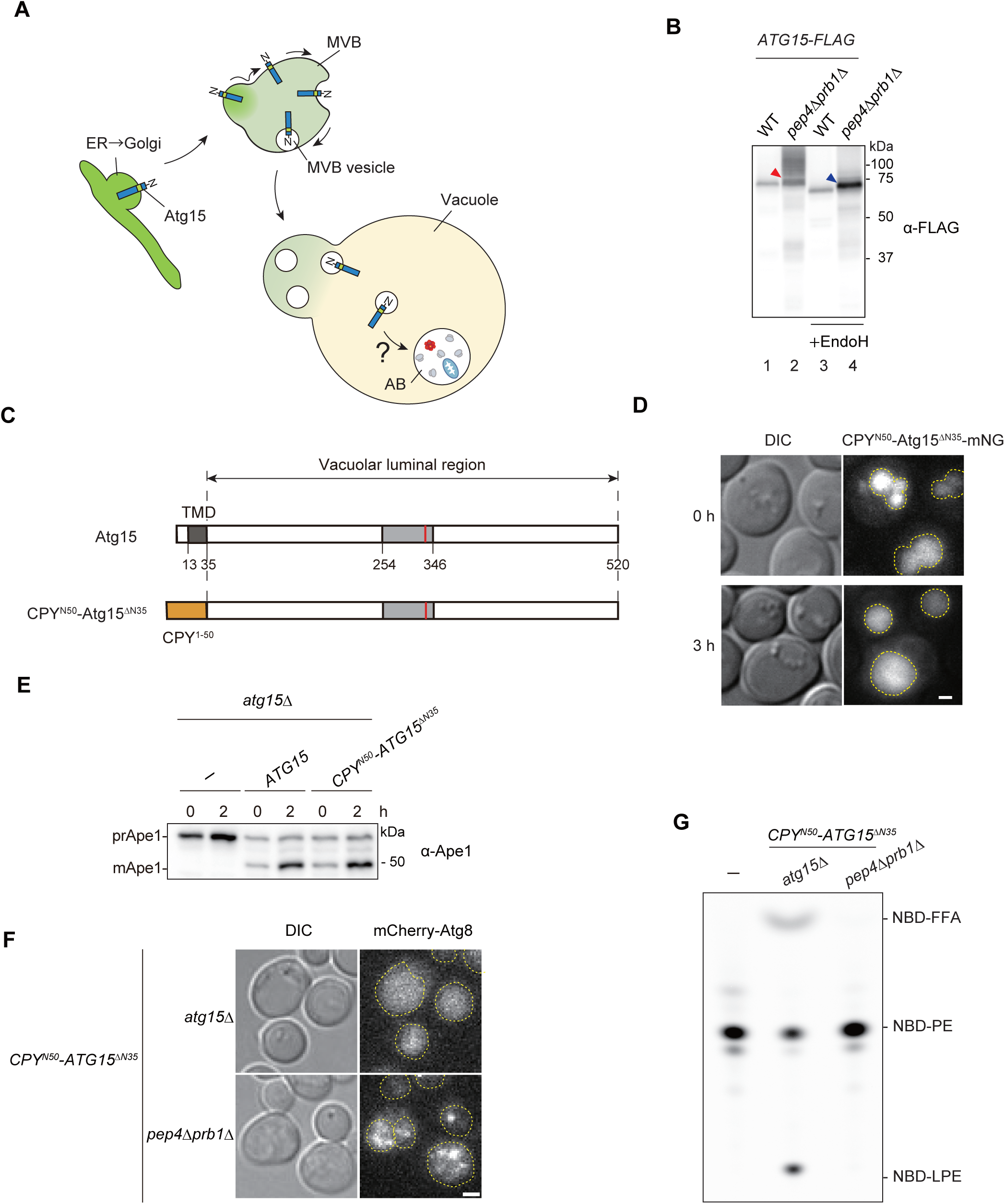
Processing is necessary for activation of the vacuolar luminal region of Atg15. (A) Schematic illustration of the transport of Atg15 from the ER/Golgi to the vacuole via the MVB pathway. (B) Western blotting of Atg15-FLAG in the vacuole. Vacuolar fractions were obtained from *atg15*Δ or *pep4*Δ *prb1*Δ cells expressing Atg15-FLAG. The samples were treated with or without Endo H and analyzed using α-FLAG and α-Pho8 antibodies. Red arrowhead, 75 kDa band of glycosylated whole Atg15-FLAG; blue arrowhead, unglycosylated whole Atg15-FLAG. The amount of the lysates used for this analysis was shown in Fig EV2C. (C) Schematic diagram of Atg15 and CPY^N50^-Atg15^ΔN35^. (D) Fluorescence microscopy image of *atg15*Δ cells expressing CPY^N50^-Atg15^ΔN35^-mNG under the control of the endogenous *ATG15* promoter from a multicopy plasmid. Cells grown in SD/CA medium were treated with rapamycin for 3 h. Scale bar, 1 μm. Dashed lines indicate vacuole boundaries. (E) Analysis of the maturation of prApe1 in *atg15*Δ cells expressing Atg15 or CPY^N50^-Atg15^ΔN35^ under the control of the endogenous *ATG15* promoter from a multicopy plasmid. Cells grown in SD/CA medium were treated with rapamycin for 2 h. prApe1 and mApe1 were detected using α-Ape1 antibodies. (F) Localization of mCherry-Atg8 in *atg15*Δ cells or *pep4*Δ *prb1*Δ cells, each expressing CPY^N50^-Atg15^ΔN35^. Cells grown in YPD medium were treated with rapamycin for 3 h. Scale bar, 1 μm. See also Movie EV1. Dashed lines indicate vacuole boundaries.(G) Lipase activities of CPY^N50^-Atg15^ΔN35^-expressing *atg15*Δ cells and *pep4*Δ *prb1*Δ cells. The amount of the lysates used for these analyses is shown in Fig EV2F.

We next tried to express Atg15 lacking its transmembrane domain to gain more insights into processing in the vacuole. Expression of Atg15 lacking residues 1-35 (“vacuolar luminal region”, Atg15^ΔN35^) in fusion with 50 N-terminal residues (1-50) of carboxypeptidase Y (CPY ^N50^) allows for the transport of the resulting CPY^N50^-Atg15^ΔN35^ (Fig 2C) to the vacuole via the CPY pathway (Johnson et al., 1987). By tagging CPY^N50^-Atg15^ΔN35^ with mNeonGreen (mNG) at its C-terminus, we confirmed that this fusion protein localizes to the vacuole under growing and autophagy-inducing conditions (Fig 2D). To determine whether CPY^N50^-Atg15^ΔN35^-expressing cells are able to disrupt ABs in the absence of endogenous Atg15, processing of precursor Ape1 (prApe1), a cargo protein of both the autophagy and Cvt pathways, was monitored. Following disruption of ABs and Cvt vesicles, prApe1 is processed to a mature form (mApe1) by vacuolar proteases (Baba et al., 1997; Klionsky et al., 1992). mApe1 was observed to occur normally in *atg15*Δ cells expressing CPY^N50^-Atg15^ΔN35^ under both growing and autophagy-inducing conditions (Fig 2E and EV2D). Therefore, we concluded that the vacuolar luminal region is sufficient to generate active Atg15 able to disrupt ABs and Cvt vesicles.

We further examined whether CPY^N50^-Atg15^ΔN35^ is able to disrupt ABs in the absence of Pep4 and Prb1. ABs were labeled with mCherry-tagged Atg8 (Kirisako et al., 1999). Upon induction of autophagy, mCherry-Atg8 signal was homogenously diffused in the vacuole of wild-type cells (Fig EV2E). *atg15*Δ cells expressing CPY^N50^-Atg15^ΔN35^ exhibited a similar pattern (Fig 2F; Movie EV1), indicating that autophagy proceeded normally in these cells. On the other hand, in *pep4*Δ *prb1*Δ cells expressing CPY^N50^-Atg15^ΔN35^ cells, intact ABs were observed in vacuoles (Fig 2F; Movie EV1). We also observed that the vacuolar lysates of CPY^N50^-Atg15^ΔN35^ expressing cells do not exhibit lipase activity in the absence of Pep4 and Prb1 (Fig 2G and Fig. EV2F). Together, these data suggest that release of Atg15 from MVB vesicles is not sufficient for acquisition of lipase activity, but rather that processing within the vacuolar luminal region is required.

### The vacuolar luminal region of Atg15 binds to the AB membrane

In order to further analyze Atg15, we sought to understand the behavior of the vacuolar luminal region of this protein. Atg15 and CPY^N50^-Atg15^ΔN35^ were tagged with mNG at their C-termini and expressed in *pep4*Δ *prb1*Δ cells to avoid cleavage in the presence of Pep4 and Prb1 (Fig 2D). Super-resolution fluorescence microscopy showed rapidly moving Atg15-mNG puncta within vacuoles (Fig 3; Movie EV2). These most likely represent MVB vesicles, as previously reported (Hirata et al., 2021). Occasionally, distinct bright foci were observed, likely to be an aggregate form. On the other hand, CPY^N50^-Atg15^ΔN35^-mNG showed ring structures surrounding mCherry-Atg8 foci (Fig 3; Movie EV3), indicating that the vacuolar luminal domain is bound to the AB membrane. Upon delivery to the vacuole, Atg15 might bind to the AB membrane once this protein detaches from the MVB.

**Figure 3.**
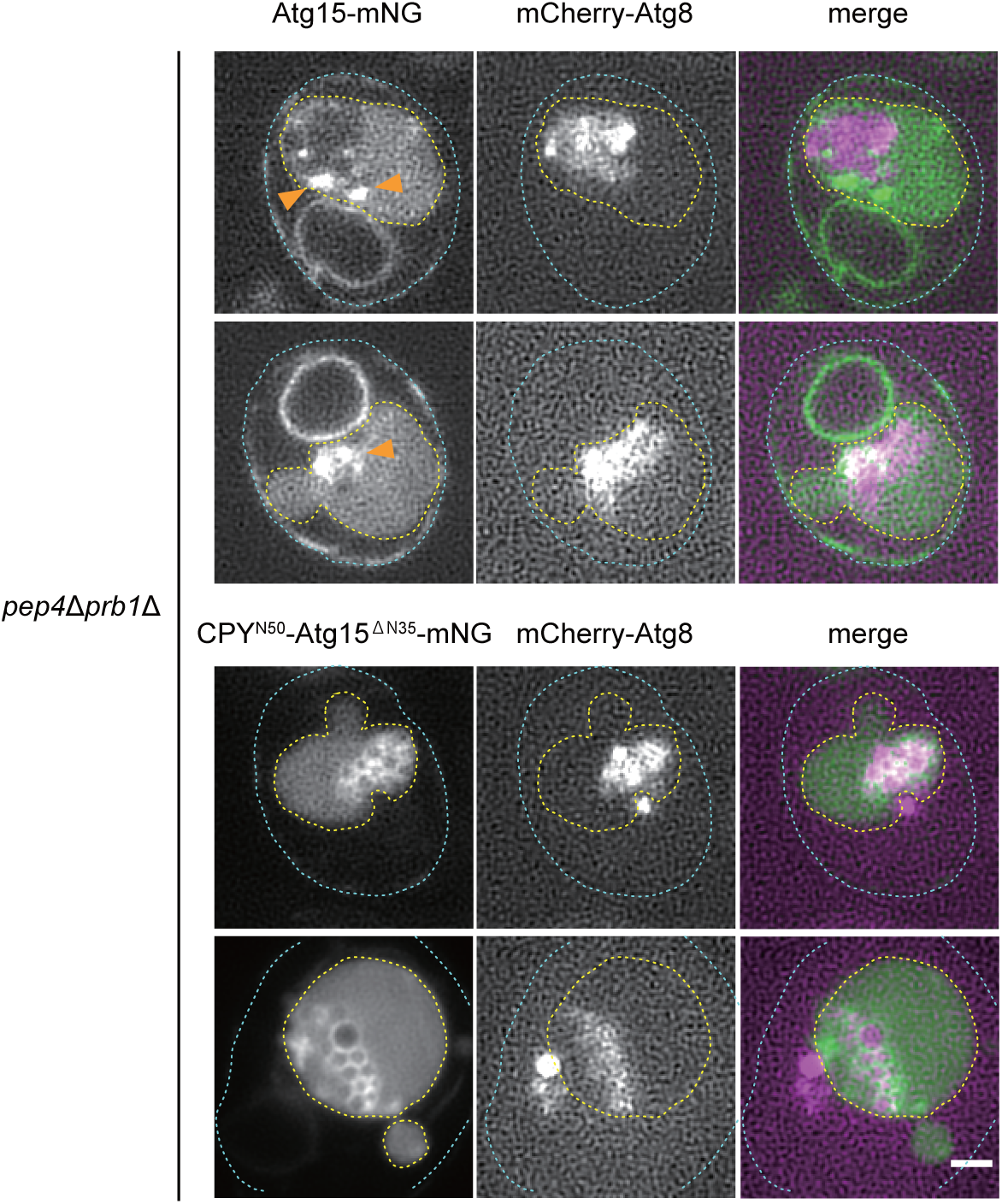
The vacuolar luminal region of Atg15 binds to the AB membrane. Super resolution microscopy image of *pep4*Δ *prb1*Δ cells expressing Atg15-mNG or CPY^N50^-Atg15^ΔN35^-mNG. Cells were grown in YPD medium and treated with rapamycin for 3 h. ABs are indicated by mCherry-Atg8 fluorescence. The orange arrowheads indicate aggregated Atg15-mNG. Dashed lines indicate vacuole (yellow) and cell (cyan) boundaries. Scale bar, 1 μm. See also Movie EV2 and 3.

### Isolation of active Atg15 from the pellet fraction of the vacuole

Due to the cleavage of the terminal tag (Fig 2B), we constructed an internally FLAG-tagged Atg15 variant to further study the processing of Atg15 in the vacuole. We first set out to identify the region necessary to disrupt ABs by making a series of N- and C-terminal truncation mutants of Atg15 that we fused with CPY^N50^. By expressing these truncation mutants in *atg15*Δ cells, we found that the deletion of residues 1-58 had little effect on the maturation of prApe1, but further deletion of Arg59 severely impaired maturation (Fig EV3A). When C-terminal residues 467-520 were deleted, prApe1 was still processed into mApe1, whereas the deletion of Trp466 caused an almost complete maturation defect (Fig EV3B). These results suggest that residues 59-466 of Atg15 are required to disrupt ABs, which is in close agreement with a previous study (Hirata et al., 2021). After several trials, we found that the FLAG tag can be inserted after Asp168 (Atg15^iFLAG^) without affecting prApe1 maturation (Fig 4A).

**Figure 4.**
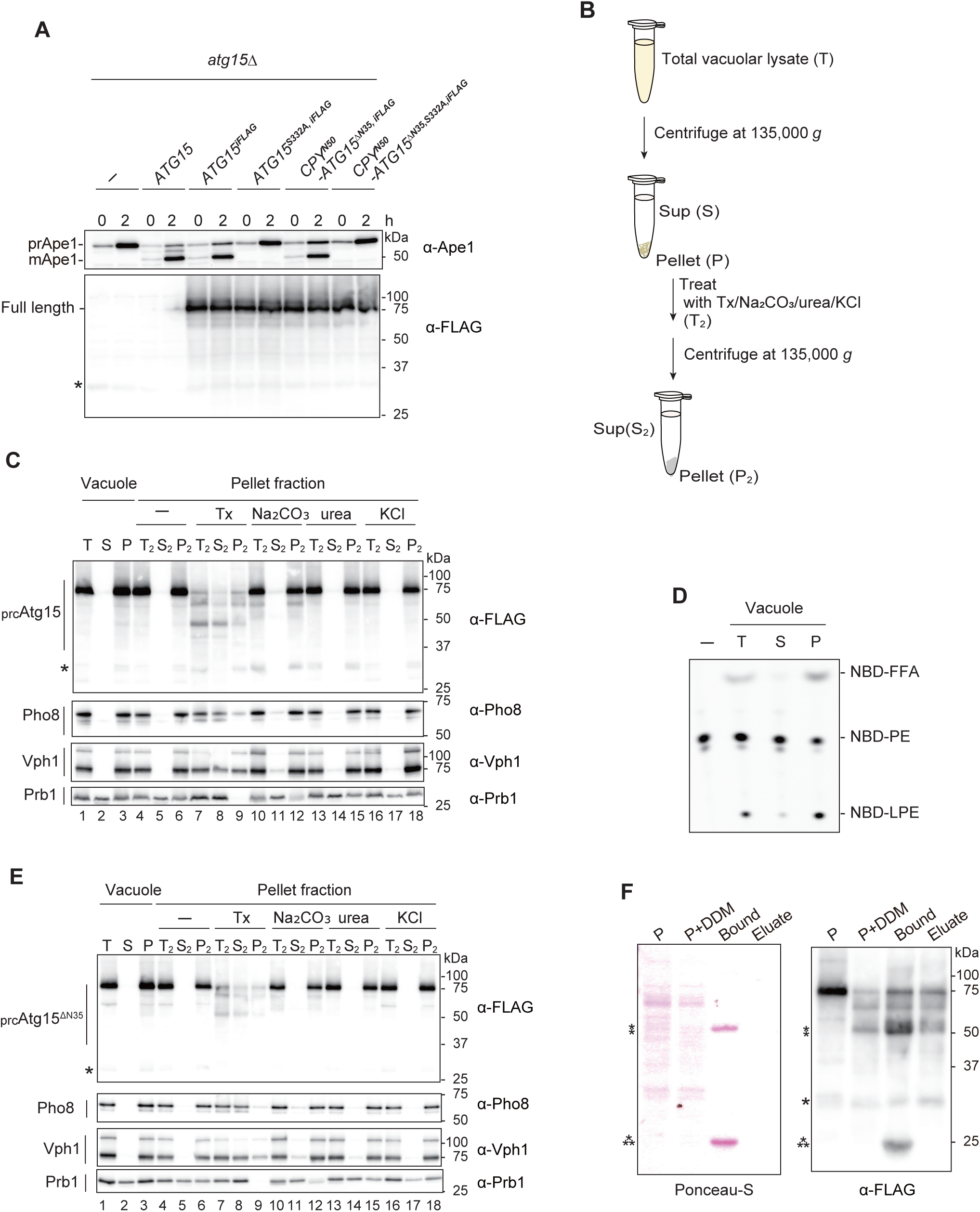
Isolation of active Atg15 from pellet fraction of the vacuole. (A) Maturation of prApe1 and expression of Atg15^iFLAG^, Atg15^S332A,iFLAG^, CPY^N50^-Atg15^ΔN35,iFLAG^ or CPY^N50^-Atg15^ΔN35,S332A,iFLAG^. Cells were grown in SD/CA medium and treated with rapamycin for 2 h. *, nonspecific band (B) Overview of the strategy employed for the fractionation of _prc_Atg15s. Total vacuolar lysate (*T*) was fractionated to a supernatant (*S*) fraction and pellet (*P*) by centrifugation. The pellet (*P* = *T_2_*) was then resuspended in buffer containing 2% Tx, 0.1 M Na_2_CO_3_, 2.5 M urea, or 1 M KCl and subjected to a second round of centrifugation to yield a pellet fraction (*P_2_*) and supernatant (*S_2_*) fraction. (C) Western blotting of _prc_Atg15 and vacuolar membrane proteins. Vacuolar lysates from Atg15^iFLAG^-expressing cells were subjected to fractionation as in (B). Equivalent amounts (relative to the starting materials) of each fraction were analyzed. (D) Lipase activities of *T*, *S*, *P* fraction (panel C lane 1-3). (E) Western blotting of the processed forms of CPY^N50^-Atg15^ΔN35,iFLAG^ (_prc_Atg15^ΔN35^s) and vacuolar membrane proteins (as in C). (F) Western blotting of _prc_Atg15s obtained by immunoprecipitation. Untreated vacuolar pellet fraction (*P*), vacuolar pellet fraction treated with 0.5% DDM (*P+DDM*), _prc_Atg15s pulled-down by α-FLAG antibody conjugated agarose beads (*Bound*), and _prc_Atg15s eluted from *Bound* by an excess amount of 3xFLAG peptides (*Eluate*) were analyzed. Samples equivalent to 0.2% of *P* and *P + DDM*, and 5% of *Bound* and *Eluate* were subjected to Ponceau-S staining (left) and western blotting using α-FLAG antibodies (right). *, nonspecific band; ⁑, mouse-IgG heavy chain; ⁂, mouse-IgG light chain.

We next undertook biochemical characterization of the processed forms of Atg15^iFLAG^ (hereafter referred as _prc_Atg15s). Vacuolar lysates from cells expressing Atg15^iFLAG^ were subjected to centrifugation at 135,000 *g* (Fig 4B). _prc_Atg15s were barely detected in the supernatant: almost all were recovered in the pellet fraction (Fig 4C lanes1-3). In addition, lipase activity was also mostly detected in the pellet fraction (Fig 4D). In order to extract processed Atg15 from pellets, we next treated the pellet fraction with a range of conditions, including high pH, urea, high salt, or detergent. High pH, urea and high salt are known to strip peripheral proteins from membranes, whereas detergent solubilizes both integral and peripheral membrane proteins. _prc_Atg15s were not extracted following high pH, urea or high salt treatments (Fig 4C, lanes 10-18), but were partially solubilized with the detergent Triton X-100 (Tx) (Fig 4C lanes 7-9). Vph1 and Pho8, which are integral vacuolar membrane proteins, were also extracted by Tx. Other detergents, such as *n*-Dodecyl-β-D-maltoside (DDM), solubilized the _prc_Atg15s to a nearly equal extent as Tx (Fig EV3C). We further examined the membrane-binding properties of CPY^N50^-Atg15^ΔN35^, finding that the vacuolar luminal region is tightly bound to the membrane (Fig 4E), and also found that _prc_Atg15s were partially degraded when subjected to detergents (Fig 4C lanes 7-9 and Fig 4E lanes 7-9, EV3C). This likely reflects an increased sensitivity of _prc_Atg15s to proteases following solubilization.

We further investigated the lipase activity of prcAtg15s, but failed to detect any such activity in the presence of detergents (Fig EV3D and E). To obtain detergent-free _prc_Atg15s, homogenate of the vacuolar pellet was first solubilized with 0.5% DDM in the presence of α-FLAG antibody conjugated agarose beads, following which the beads were collected and subsequently washed with a buffer solution to remove unbound materials and the detergent. The beads were then incubated with an excess amount of 3xFLAG peptide to elute _prc_Atg15s. Although the amount of the final eluted fraction was too small to detect by ponceau staining, several processed forms of Atg15^iFLAG^ were detected by western blotting with α-FLAG antibodies (Fig 4F).

### The active Atg15 is phospholipase B

Following the above procedures, we prepared the eluate of both _prc_Atg15s and _prc_Atg15^S332A^ s (Fig 5A) and characterized their lipase activities. By incubation with _prc_Atg15 eluates, NBD-PE levels as determined by TLC decreased in a time-dependent manner to an almost undetectable level at 6 h. NBD-LPE increased up to 1 h, and then decreased. NBD-FFA increased throughout the incubation, indicating that NBD-FFA is generated from both NBD-PE and NBD-LPE (Fig 5B). In contrast, the _prc_Atg15^S332A^ eluate did not exhibit any lipase activity at all (Fig 5B).

**Figure 5.**
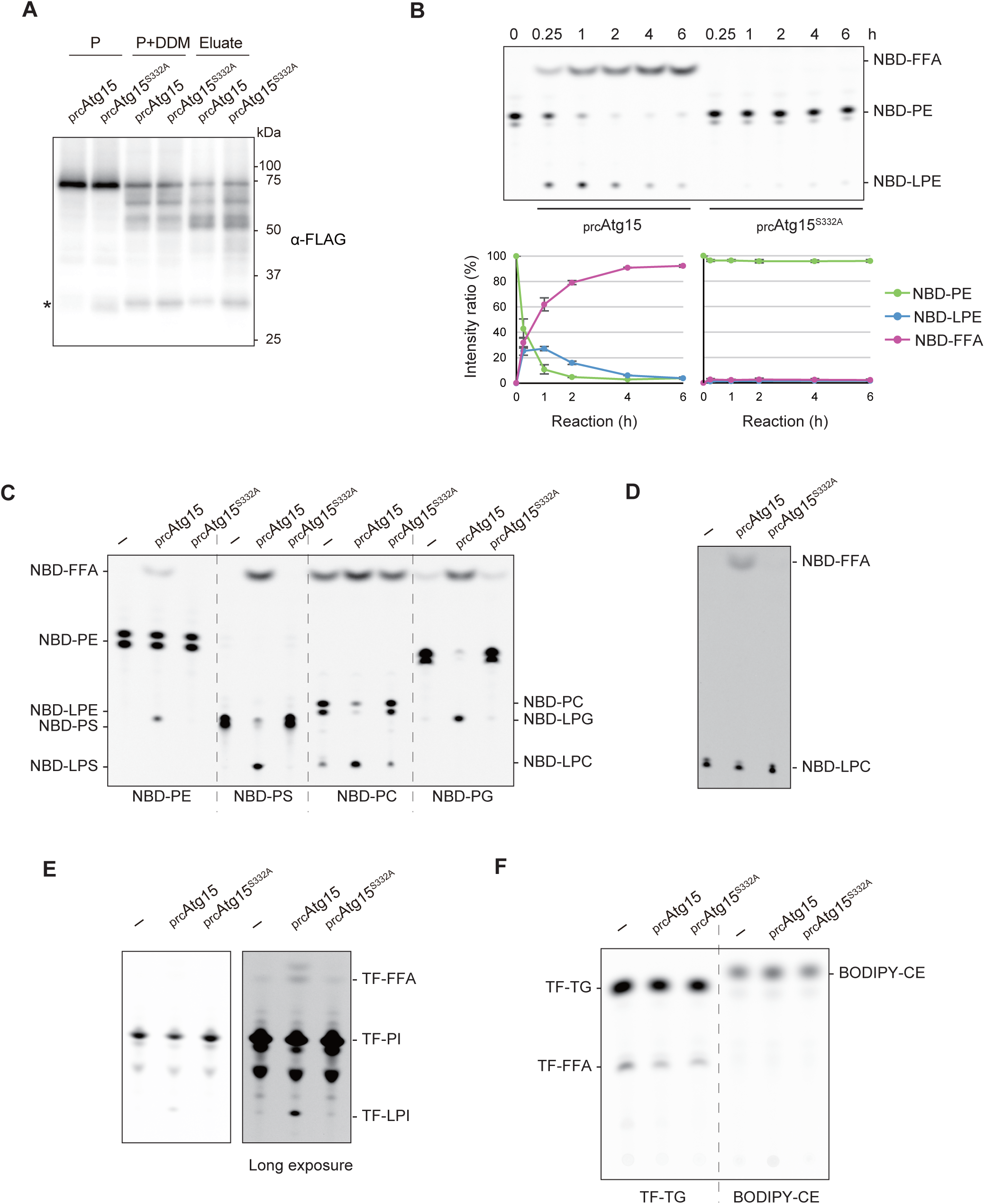
Activated Atg15 is a phospholipase B of broad head-group specificity. (A) The eluates of _prc_Atg15s and _prc_Atg15^S332A^s were obtained as in Fig 4F. Samples equivalent to 0.3% of *P* and *P+DDM,* and 22 % of *Eluate* were subjected to western blotting with α-FLAG antibodies. *, nonspecific band. (B) Time course of lipase activity in _prc_Atg15s and _prc_Atg15^S332A^s. At each point, 3 μl of *Eluates* in (A) were incubated with NBD-PE. Representative results from three independent experiments are shown (upper panel). Intensity ratios (lower panel) were calculated as follows: intensity ratio (%) = (peak area for NBD-PE, NBD-LPE, or NBD-FFA) x100 / (peak area for NBD-PE + peak area for NBD-LPE + peak area for NBD-FFA). Error bars represent means ± S.D. of three independent experiments. (C-D) Lipase activity toward NBD-PE, NBD-PS (18:1-12:0), NBD-PC (18:1-12:0), and NBD-PG (18:1-12:0), each labeled at *sn-2* (C, E), and NBD-LPC (12:0) (D). Each phospholipid was incubated with 1 μl of *Eluate* (A) for 1 h. (E) Lipase activity toward TF-PI (18:1-6:0) labeled at *sn-2.* TF-PI was incubated with 4 μl of *Eluate* (A) for 2 h. (F) Lipase activity toward TF-TG and BODIPY-CE. Each non-polar lipid was incubated with 4 μl of *Eluate* (A) for 2 h.

Next, we examined substrate specificity. NBD-PS was hydrolyzed as NBD-PE (Fig 5C). Commercially available NBD-PC and NBD-PG had been somewhat decomposed, but were clearly hydrolyzed by _prc_Atg15 eluates (Fig 5C). Meanwhile, NBD-labeled lysophosphatidylcholine (LPC) was cleaved to NBD-FFA (Fig 5D), and we also observed hydrolysis of topflour-labeled PI (TF-PI) to TF-lysophosphatidylinositol (TF-LPI) and TF-FFA (Fig 5E). On the other hand, we were unable to detect any activity toward non-polar lipids such as TG and cholesterol ester (CE) in our assay conditions (Fig 5F). These results correspond well with those from vacuolar lysates (Fig EV4). Taken together, these data indicate that active Atg15 is a phospholipase B of broad head-group specificity.

### The active form of Atg15 is sufficient to disrupt ABs

We next set out to develop an in vitro assay system to evaluate AB disruption (Fig 6A). To this end, we employed our recently established method for the isolation of ABs that retain autophagosome cargo (Kawamata et al., 2022), which allowed us to use the purified ABs as a native substrate of Atg15. We isolated ABs from *atg15*Δ cells expressing GST-GFP, which localizes to the cytosol and is delivered to the vacuole via autophagy. prApe1 and GST-GFP in the AB were resistant to ProK treatment (Fig 6B, lanes 1 and 2). Upon treatment of ABs with detergent, prApe1 and GFP-GST were cleaved to produce degraded Ape1 (dApe1) and free GFP (Fig 6B lane 3), respectively. Therefore, the amount of dApe1 or free GFP reflects the degree of AB membrane disruption. When ABs were treated with _prc_Atg15 eluate, dApe1 and free GFP appeared (Fig 6B lane 4 and 6), indicating that _prc_Atg15 eluates are capable of the disruption of AB membranes. Furthermore, the extent of this disruption was dependent on the amount of _prc_Atg15 eluate added (Fig 6B, lanes 4 and 6). On the other hand, treatment of ABs with _prc_Atg15^S332A^ did not result in membrane disruption (Fig 6B, lanes 5 and 7). These results indicate that active Atg15 is sufficient to disrupt AB membranes in vitro.

**Figure 6.**
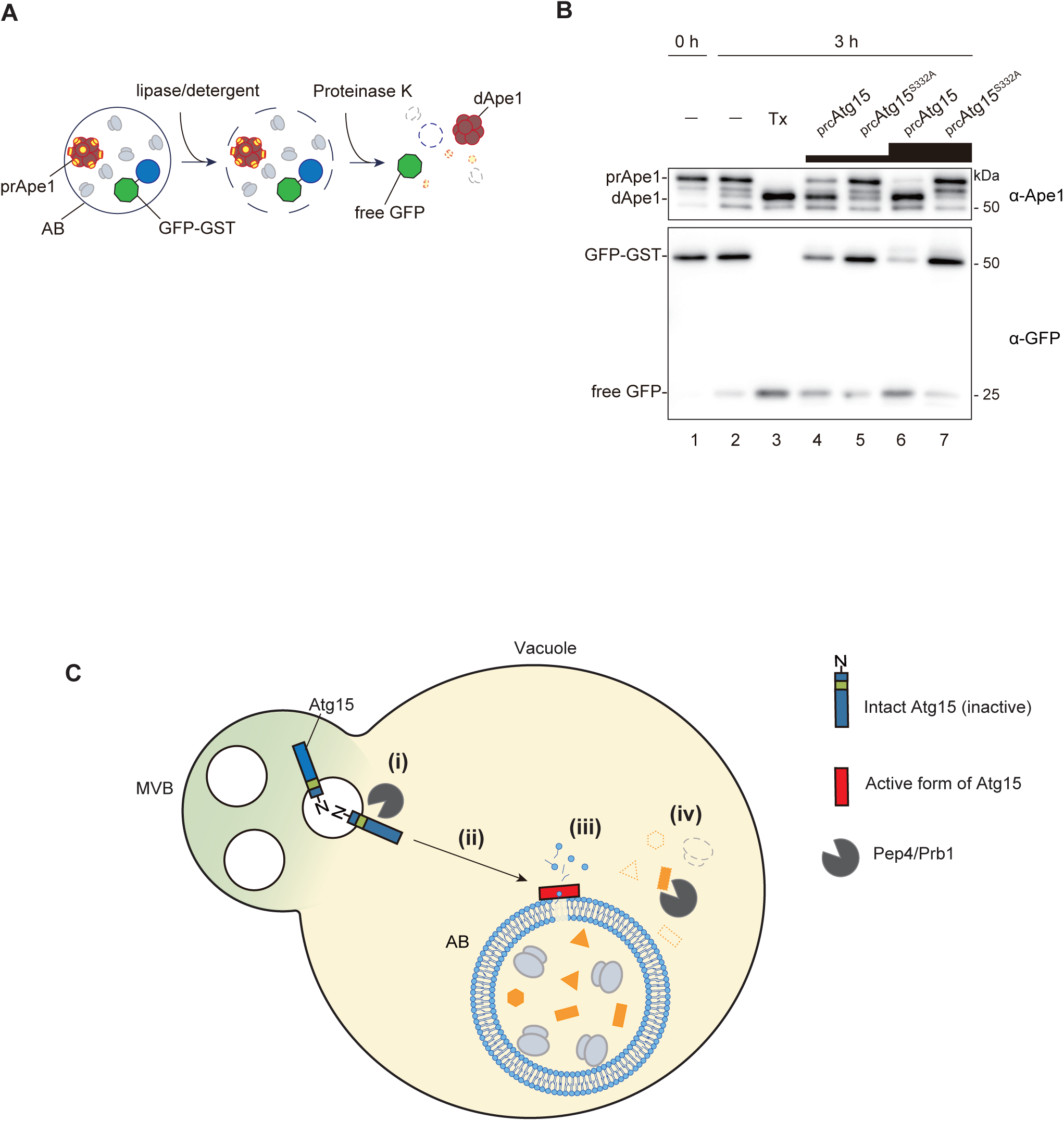
Active Atg15 is sufficient to disrupt Abs. (A) Schematic view of the in vitro assay employed to evaluate AB disruption. (B) Disruption of ABs by active Atg15. Isolated ABs were incubated with 0.2% Tx, _prc_Atg15 or _prc_Atg15^S332A^ eluates at 30 °C for 0 or 3 h. Each eluate was added at a volume ratio of 1:3. The reaction mixture was subjected to western blotting using α-Ape1 and α-GFP antibodies, respectively. (C) A model of AB disruption by active Atg15 in the vacuole. (i) Upon delivery to the vacuole, Atg15 is cleaved from MVB vesicles by Pep4/Prb1. (ii) Processed forms of Atg15 are tightly bound to the AB membrane. Atg15 is activated by Pep4/Prb1 during or following localization to the AB membrane. (iii) The active form of Atg15 hydrolyzes acyl ester linkages of phospholipids of AB membranes, releasing two acyl groups and disrupting the membrane. (iv) AB contents are released into the vacuole, where they are degraded by vacuolar hydrolases including Pep4 and Prb1.

Overall, we propose a model for the processing of Atg15 to an active form and the subsequent lysis of AB membranes in the vacuole by active Atg15 (Fig 6C). First, Atg15 is delivered to the vacuole by the MVB pathway, where it is immediately cleaved from MVB vesicles by Pep4/Prb1. The processed forms are tightly bound to the AB membrane and activated by Pep4/Prb1 during localization to or once physically associated with the AB membrane. Following activation, Atg15 is able to hydrolyze ester bonds at the *sn*-*1* and *sn*-*2* positions of AB membrane phospholipids. Finally, released AB contents are degraded by vacuolar hydrolases including Pep4 and Prb1, completing the degradation of autophagy cargos.

## Discussion

Both Atg15 and the proteases Pep4/Prb1 are necessary for AB disruption, but how these proteins are involved in this process has remained unclear. In this study, we established an in vitro lipase assay for Atg15-derived lipase activity, finding that Pep4 and Prb1 process Atg15 to an activated form that exhibits lipase activity. Furthermore, we used isolated ABs to develop an in vitro system allowing for the evaluation of AB membrane disruption which showed that activated Atg15, which has phospholipase B activity, is sufficient to disrupt AB membranes.

### Activation of Atg15 by Pep4/Prb1-mediated processing

Our in vitro lipase assay helped clarify the relationship between Atg15 lipase activity and other vacuolar proteins that have previously been implicated in AB disruption. We showed that appearance of Atg15 lipase activity completely depends on Pep4 and Prb1 (Fig 1F). The fact that cells overexpressing Atg15 show no lipase activity in the absence of Pep4/Prb1 (Fig EV1E) strongly suggests that Pep4/Prb1 activate Atg15 by direct processing of the protein rather than indirectly via the degradation of an inhibitor of Atg15.

Our analyses suggest that Atg15 is cleaved at the region adjacent to the transmembrane domain by Pep4/Prb1 to liberate it from MVB vesicles, after which it binds AB membranes (Fig 2B and Fig 3). We also found that the processing of the vacuolar luminal region of Atg15 is necessary for Atg15 to acquire lipase activity (Fig 2G). Creation of an internally FLAG-tagged Atg15 enabled us to analyze the processing of Atg15 by immunopurification (Fig 4). Using this strategy, we successfully obtained active form(s) of Atg15 with lipase activity (Fig 4F). The 3D structure of Atg15, which is currently unsolved, will provide important clues that allow for a more nuanced understanding of the activation mechanism.

### Enzymatic features of Atg15 following activation

Ramya & Rajasekharan previously reported that Atg15 purified from microsomal membrane fractions of yeast preferentially hydorolyzes PS rather than other phospholipids (Ramya & Rajasekharan, 2016). In our study, active Atg15 did not show such a preference for PS: PC, PG, and PE were also hydrolyzed (Fig 5C). This discrepancy may be related to technical differences in assay conditions: our lipase assay was performed at physiological pH, whereas Ramya & Rajasekharan measured at pH 8.0.

Importantly, our results reveal for the first time that active Atg15 exhibits phospholipase B activity (Fig 5B). The ability of phospholipase B enzymes to cleave both acyl groups of phospholipids is likely important for efficient membrane disruption, and the low substrate specificity of Atg15 may also be advantageous for the degradation of diverse membrane lipids of Cvt bodies, MVB and microautophagic vesicles, as well as organellar membranes such as mitochondria, ER, peroxisome following their delivery to the vacuole. For non-polar lipids (TG and CE), we were unable to observe hydrolysis by activated Atg15 using our lipase assay (Fig 5F). Further analysis is needed to determine if Atg15 has such activity under different conditions, or if there are alternative lipases in the vacuole that are able to hydrolyze such non-polar lipids.

### Hydrophobic features of Atg15

Many of the vacuolar hydrolytic enzymes undergo processing by proteases (Hecht et al., 2014). We assume that Atg15 is maintained in an inactive state during trafficking from the ER to Golgi, instead becoming active only after delivery to the vacuole. Atg15 is very sensitive to proteases, especially when detached from membranes in the vacuole (Fig 4C, E, F). Free Atg15 is therefore rapidly degraded in the vacuole, as previously reported (Epple et al., 2001; Teter et al., 2001). These properties may be critical for the spaciotemporal regulation of potentially deleterious lipase activity.

Our analysis of the vacuolar luminal region raises two intriguing implications. First, this region binds to the AB membrane, but not the vacuolar membrane (Fig 3). This finding likely explains why Atg15 is unable to digest the vacuolar membrane. We also found that this region binds the AB membrane even when the protein is not activated (Fig 2F, G, Fig 3), suggesting that activation of Atg15 might occur on AB membranes, which provides an excellent way to restrict Atg15 lipase activity to specific membranes. The apparent preference of Atg15 for AB membranes may be explained by the high membrane curvature of these membranes (the vacuolar membrane is generally characterized by very low or negative curvature). Alternatively, glycoproteins abundant on inner surface of the vacuolar membrane may not allow Atg15 to bind to the vacuolar membrane.

The second implication of our study is that the processed forms of the vacuolar luminal region of Atg15 are tightly bound to the membrane in a manner reminiscent of a transmembrane protein (Fig 4C and E). Although a search of the domain database (Letunic et al., 2020) revealed no identified transmembrane domains within the vacuolar luminal region (Fig 1A), Atg15 probably contains membrane-binding sites besides the active center, as observed in many other lipases (Cao et al., 2013). Indeed, the structure of Atg15, as predicted by AlphaFold2 (Tunyasuvunakool et al., 2021), shows that Atg15 has several clusters of hydrophobic residues facing toward the exterior of the protein, some of which may become embedded in AB membranes.

### Physiological functions of the vacuole/lysosome lipases

Recently, lysosomal phospholipase A2 (PLA2G15/LPLA-2) was identified as a key lipase responsible for degrading membranes in lysosomes. Deletion of PLA2G15/LPLA-2 results in the formation of enlarged lysosomes that accumulate abundant membranous structures (Li et al., 2021). However, no studies have yet assessed how PLAG15/LPLA-2 in lysosomes degrade membrane lipids delivered via autophagy or other transport routes. The complexity of lysosomal hydrolases may make it difficult to analyze the protein network responsible for membrane disruption within lysosomes. The relative simplicity of yeast cells was particularly advantageous in our study, allowing us to clarify the relationship between proteases and lipase activity in the vacuole.

Characterization of lipase activity in the vacuole/lysosome is essential to understand how lipids are recycled. Our identification of Atg15 as a phospholipase B with broad substrate specificity indicates that various species of FFAs, LPLs, and water soluble glycerophosphodiesters (GPDs) are generated from membranes implicated in autophagy.

Very recently, Spns1 was identified as a transporter that effluxes LPC and LPE in the lysosome for their recycling into cellular phospholipid pools (He et al., 2022). Further, a recent study Laqtom et al. showed that CLN3, which is a lysosomal transmembrane protein involved in Batten disease, is required for the efflux of GPDs from lysosomes in mouse (Laqtom et al., 2022). This protein shares the same function with its yeast homolog, Btn1 (Mirza et al., 2019). Considering these reports, LPLs and GPDs in the vacuole might be reused as lipids or other components. On the other hand, excessive accumulation of FFAs in the vacuole/lysosome could lead to lipotoxicity and vacuole/lysosome dysfunction. Further studies on the fate of these degradation products will uncover the impact of lipid degradation by autophagy on cellular metabolism.

## Materials and Methods

### Lipids used in this study

NBD-PE (1-oleoyl-2-{12-[(7-nitro-2-1,3-benzoxadiazol-4-yl)amino]dodecanoyl}-*sn*-glycero-3- phosphoethanolamine), NBD-PS (1-oleoyl-2-{12-[(7-nitro-2-1,3-benzoxadiazol-4- yl)amino]dodecanoyl}-*sn*-glycero-3-phosphoserine), NBD-PC (1-Oleoyl-2-[12-[(7-nitro-2-1,3- benzoxadiazol-4-yl)amino]dodecanoyl]-sn-Glycero-3-Phosphocholine), NBD-PG (1-oleoyl-2- {12-[(7-nitro-2-1,3-benzoxadiazol-4-yl)amino]dodecanoyl}-sn-glycero-3-[phospho-rac-(1- glycerol)]), TF-PI (1-oleoyl-2-{6-[4-(dipyrrometheneboron difluoride)butanoyl]aminhexanoyl- *sn*-glycero-3-phosphoinositol), NBD-LPC (1-{12-[(7-nitro-2-1,3-benzoxadiazol-4- yl)amino]dodecanoyl}-2-hydroxy-sn-glycero-3-phosphocholine), and TF-TG (1,2-dioleoyl-3- [11-(dipyrrometheneboron difluoride)undecanoyl]-sn-glycerol) were obtained from Avanti Polar Lipids. BODIPY-CE (cholesteryl 4,4-difluoro-5,7-dimethyl-4-bora-3a,4a-diaza-*s*-indacene-3- dodecanoate) was obtained from Life Technologies. Lipids (with the exception of NBD-LPC) were dissolved in 5 mM CHAPS at a concentration of 125 ng/μl and sonicated for 1-5 min. NBD-LPC was dissolved in water.

### Buffers used in this study

Buffer A (10 mM MES-Tris pH 6.9, 0.2 M sorbitol, 12% w/v Ficoll 400, 0.1 mM MgCl_2_), buffer B (10 mM MES-Tris pH 6.9, 0.2 M sorbitol, 8% w/v Ficoll 400, 0.1 mM MgCl_2_,), buffer B’ (10 mM MES-Tris pH 6.9, 0.2 M sorbitol, 4% w/v Ficoll 400, 0.1 mM MgCl_2_), buffer C (10 mM MES-Tris pH 6.9, 0.2 M sorbitol, 0.1 mM MgCl_2_), buffer D (10 mM MES-Tris pH 6.9, 5 mM MgCl_2_, 150 mM KCl), buffer E (30 mM MES-Tris pH 6.9, 200 mM KCl), buffer F (30 mM MES-Tris pH6.9, 100 μM CaCl_2_, 200 mM KCl), and spheroplast buffer (50 mM Tris-HCl pH 7.5, 1.2 M sorbitol) were used.

### Yeast Strains and media

The yeast strains used in this study are listed in Table 1. Gene deletion and tagging were performed using standard PCR-based methods, as described previously (Janke et al., 2004; Knop et al., 1999), and validated by PCR. Cells were cultured at 30°C in YPD medium (1% yeast extract (Gibco), 2% Bacto peptone (Gibco), 2% glucose) or in a synthetic defined medium comprising 0.17% yeast nitrogen base without amino acids and ammonium sulfate (BD), 0.5% ammonium sulfate, 0.5% casamino acids (BD), 2% glucose, 20 μg/ml adenine sulfate, and 20 μg/ml tryptophan (SD/CA medium). To induce autophagy, cells were treated with rapamycin (LC Laboratories, R-5000) at a concentration of 0.2 μM.

**Table 1.** Yeast strains used in this study

| No. | Strain | Genotype | Source | Figures |
| --- | --- | --- | --- | --- |
| 1 | X2180-1B | *MATa SUC2 mal mel gal2 CUP1* | Yeast Genetic Stock Center | Fig 1C, F |
| 2 | MMY209 | X2180-1B; *atg15Δ::kanMX6* | this study | Fig 1C |
| 3 | KYY070 | X2180-1B; *atg15Δ::kanMX6, ura3Δ::hphNT1*<br>pRS426 | this study | Fig 2E, 4A |
| 4 | KYY127 | X2180-1B; *atg15Δ::kanMX6, ura3Δ::hphNT1*<br>pRS426-CPY(1-50)-ATG15(Δ1-35)-mNeonGreen | this study | Fig 2D |
| 5 | KYY147 | X2180-1B; *pep4Δ::kanMX6, prb1Δ::hphNT1, leu2::ATG8pro-mCherry-ATG8(G116)-zeoNT3, ura3Δ::natNT2*<br>pRS426-CPY(1-50)-ATG15(Δ1-35)-mNeonGreen | this study | Fig 3 |
| 6 | KYY185 | X2180-1B; *pep4Δ::kanMX6, prb1Δ::hphNT1, leu2::ATG8pro-mCherry-ATG8(G116)-zeoNT3, ura3Δ::natNT2*<br>pRS426-CPY(1-50)-ATG15(Δ1-35) | this study | Fig 2F, G |
| 7 | KYY194 | X2180-1B; *atg15Δ::kanMX6, ura3Δ::hphNT1*<br>pRS426-CPY(1-50)-ATG15(Δ1-35) | this study | Fig 2E |
| 8 | KYY201 | X2180-1B; *atg15Δ::kanMX6, ura3Δ::hphNT1, leu2::ATG8pro-mCherry-ATG8(G116)-zeoNT3*<br>pRS426-CPY(1-50)-ATG15(Δ1-35) | this study | Fig 2F, G |
| 9 | KYY235 | X2180-1B; *atg15Δ::kanMX6, ura3Δ::hphNT1*<br>pRS426-ATG15(168-3xFLAG-169) | this study | Fig 4A |
| 10 | KYY236 | X2180-1B; *atg15Δ::kanMX6, ura3Δ::hphNT1*<br>pRS426-ATG15(S332A, 168-3xFLAG-169) | this study | Fig 4A |
| 11 | KYY237 | X2180-1B; *atg15Δ::kanMX6, ura3Δ::hphNT1*<br>pRS426-CPY(1-50)-ATG15(Δ1-35, 168-3xFLAG-169) | this study | Fig 4A |
| 12 | KYY238 | X2180-1B; *atg15Δ::kanMX6, ura3Δ::hphNT1*<br>pRS426-CPY(1-50)-ATG15(S332A, Δ1-35, 168-3xFLAG-169) | this study | Fig 4A |
| 13 | KYY249 | X2180-1B; *pep4Δ::kanMX6, prb1Δ::hphNT1, leu2::ATG8pro-mCherry-ATG8(G116)-zeoNT3, ura3Δ::natNT2*<br>pRS426-ATG15-mNeonGreen | this study | Fig 3 |
| 14 | SMY96 | X2180-1B; *ura3Δ::hphNT1*, *atg15Δ::kanMX6*<br>pRS426-*ATG15* | this study | Fig 1C, D, E,<br>Fig 2E, Fig<br>4A |
| 15 | SMY97 | X2180-1B; *ura3Δ::hphNT1*, *atg15Δ::kanMX6*<br>pRS426-*ATG15(S332A)* | this study | Fig 1E |
| 16 | SMY260 | X2180-1B; *ura3Δ::hphNT1*, *atg15Δ::kanMX6*<br>pRS316-*ATG15* | this study | Fig 1C |
| 17 | SMY311 | X2180-1B; *pep4Δ::kanMX6*, *prb1Δ::hphNT1* | this study | Fig 1F |
| 18 | SMY345 | X2180-1B; *atg15Δ::kanMX6*, *ura3Δ::hphNT1*<br>p416-*GPD-GST-GFPhy* | (Kawamata<br>et al., 2022) | Fig 6B |
| 19 | SMY476 | X2180-1B; *atg15Δ::kanMX6*, *ura3Δ::hphNT1*, *ape1Δ::natNT2*<br>pRS426-*ATG15-3xFLAG* | this study | Fig 2B |
| 20 | SMY485 | X2180-1B; *pep4Δ::hphNT1*, *prb1Δ::kanMX6*, *ura3Δ::natNT2*,<br>*ape1Δ::zeoNT3*<br>pRS426-*ATG15-3xFLAG* | this study | Fig 2B |
| 21 | SMY491 | X2180-1B; *atg15Δ::kanMX6*, *ura3Δ::hphNT1*, *ape1Δ::natNT2*<br>pRS426-*ATG15(168-3xFLAG-169)* | this study | Fig 4C, D, F<br>Fig 5, Fig 6B |
| 22 | SMY492 | X2180-1B; *atg15Δ::kanMX6*, *ura3Δ::hphNT1*, *ape1Δ::natNT2*<br>pRS426-*ATG15(S332A, 168-3xFLAG-169)* | this study | Fig 5, Fig 6B |
| 23 | SMY493 | X2180-1B; *atg15Δ::kanMX6*, *ura3Δ::hphNT1*, *ape1Δ::natNT2*<br>pRS426-*CPY(1-50)-ATG15(Δ1-35, 168-3xFLAG-169)* | this study | Fig 4E |
| 24 | ScYO517 | X2180-1B; *prb1Δ::hphNT1* | this study | Fig 1F |
| 25 | MMY110 | X2180-1B; *pep4Δ::hphNT1* | this study | Fig 1F |

### Plasmid construction

Plasmids and primers used in this study are listed in Table 2 and 3, respectively. To generate pRS316-*ATG15* (TPL000) and pRS426-*ATG15* (TPL001), a DNA fragment containing the *ATG15* gene from 1000 nt upstream of the initiation codon was amplified by PCR from genomic DNA. The DNA fragment was inserted into pRS316 or pRS426 using *BamH*I and *Hind*III sites. To generate pRS426-*ATG15(S332A)* (TPL002), the Atg15 active site S332 (serine at amino-acid position 332) was replaced with alanine in pRS426/*ATG15* (TPL001) by inverse PCR. To generate pRS426-*ATG15*-*mNeonGreen* (YPL046), the recognition sequence of *Nhe***I** was inserted into the region upstream of the stop codon of *ATG15* in pRS426-*ATG15* (TPL001). The resulting plasmid was cut using *Nhe*I, into which a PCR-amplified DNA fragment containing the *mNeonGreen* gene (Shaner et al., 2013) was inserted using the in-Fusion reaction (Clontech). To generate pRS426-*CPY(1-50)-ATG15(Δ1-35)-mNeonGreen* (YPL073), a DNA fragment containing the *PRC1* gene was first amplified by PCR from genomic DNA and inserted into pBluescript SK(+) using *Hind*III and *EcoR*I sites. This was subsequently used as a PCR template to amplify a DNA fragment encoding the first fifty amino acids of the CPY protein, while a DNA fragment encoding the 36-520 amino acids of Atg15 fused with mNeonGreen at the C-terminus was amplified by PCR from pRS426-*ATG15*-*mNeonGreen* (YPL046). Then these DNA fragments were fused by in-Fusion reaction. To generate pRS426-*CPY(1-50)-ATG15(Δ1-35)* (YPL103), both pRS426-*CPY(1-50)-ATG15(Δ1-35)-mNeonGreen* (YPL073) and pRS426*-ATG15* (TPL001) were cut using *EcoR*I and *Hind*III, and a fusion construct containing nucleotides of CPY (1-50) and Atg15^ΔN35^ was generated by ligation (Clontech). To generate pRS426-*ATG15-3xFLAG* (TPL003), a DNA fragment encoding a glycine linker followed by a 3xFLAG sequence was inserted between the *ATG15* gene and its stop codon by inverse PCR.

**Table 2.**
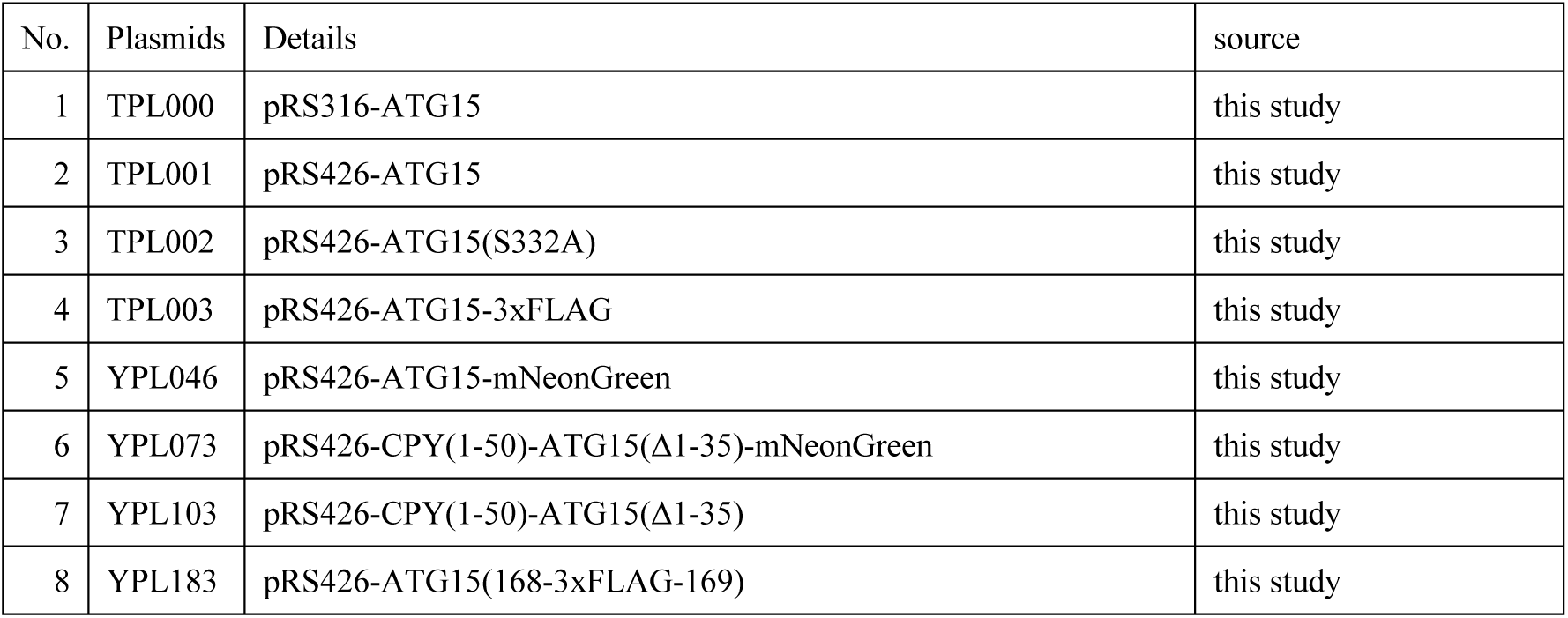
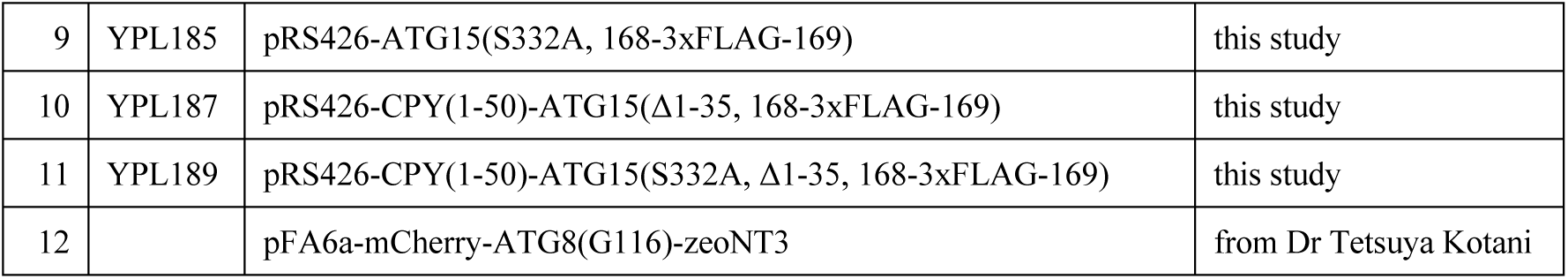
Plasmids used in this study

| No. | Plasmids | Details | source |
| --- | --- | --- | --- |
| 1 | TPL000 | pRS316-ATG15 | this study |
| 2 | TPL001 | pRS426-ATG15 | this study |
| 3 | TPL002 | pRS426-ATG15(S332A) | this study |
| 4 | TPL003 | pRS426-ATG15-3xFLAG | this study |
| 5 | YPL046 | pRS426-ATG15-mNeonGreen | this study |
| 6 | YPL073 | pRS426-CPY(1-50)-ATG15(Δ1-35)-mNeonGreen | this study |
| 7 | YPL103 | pRS426-CPY(1-50)-ATG15(Δ1-35) | this study |
| 8 | YPL183 | pRS426-ATG15(168-3xFLAG-169) | this study |
| 9 | YPL185 | pRS426-ATG15(S332A, 168-3xFLAG-169) | this study |
| 10 | YPL187 | pRS426-CPY(1-50)-ATG15(Δ1-35, 168-3xFLAG-169) | this study |
| 11 | YPL189 | pRS426-CPY(1-50)-ATG15(S332A, Δ1-35, 168-3xFLAG-169) | this study |
| 12 |  | pFA6a-mCherry-ATG8(G116)-zeoNT3 | from Dr Tetsuya Kotani |

**Table 3.**
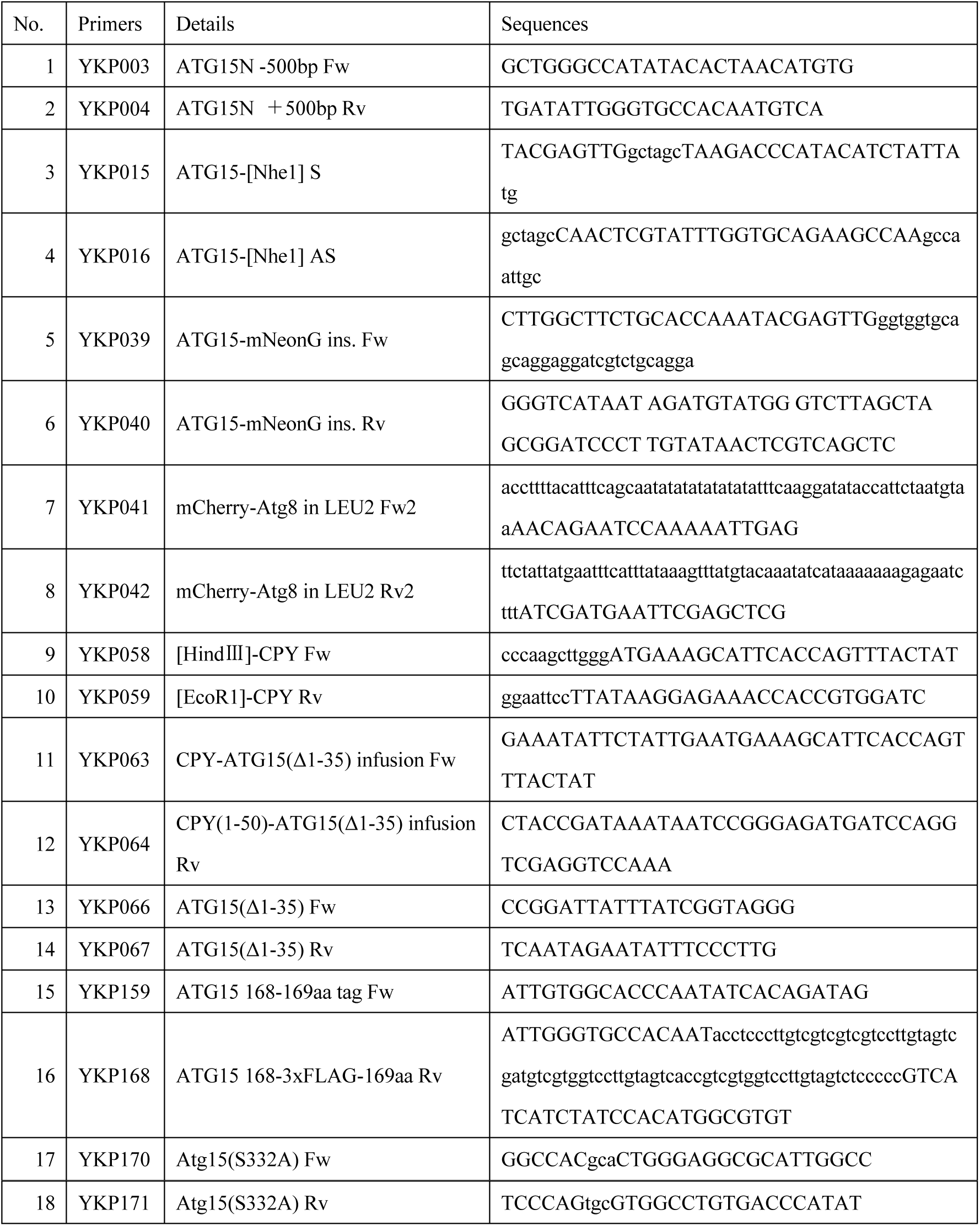
Primers used in this study

To generate the plasmids pRS426-*ATG15(168-3xFLAG-169)*(YPL183), pRS426-*ATG15(S332A, 168-3xFLAG-169)*(YPL185), pRS426-*CPY(1-50)-ATG15(Δ1-35, 168-3xFLAG-169)*(YPL187), and pRS426-*CPY(1-50)-ATG15(S332A, Δ1-35, 168-3xFLAG-169)*(YPL189), a DNA fragment encoding a glycine linker followed by a 3xFLAG sequence was inserted at the aspartic acid 168 of Atg15 by inverse PCR.

### Isolation of vacuoles

Vacuoles were obtained from cells as previously described with some modifications (Kawamata et al., 2022; Ohsumi and Anraku, 1981; Takeshige et al., 1992). Cells grown in 10 L YPD to OD600 =1.0-2.0 were treated with 0.2 μM rapamycin for 1 h. Cells were then collected and washed once with water. Spheroplasts were prepared by incubation of cells with Zymolyase 100T (Nacalai Tesque) at 10 μg/OD_600_ unit in 600 ml spheroplast buffer for 30 min. Spheroplasts were then collected by centrifugation, washed with spheroplast buffer, resuspended in 130 ml of ice-cold buffer A and homogenized using a Dounce homogenizer on ice. The lysate was then divided into six volumes that were each transferred to a new ultracentrifuge tube, and layered with 10 ml of buffer B. These layered samples were centrifuged at 72,000 *g* in a P28S swing rotor (Hitachi Koki) for 30 min at 4°C. The top, white layer of each tube was collected and combined into a new ultracentrifuge tube. This crude vacuole isolate was then mixed with 9 ml of buffer B, divided into two volumes that were each transferred to a new centrifuge tube, overlaid with 13 ml of buffer B’ and 13 ml of buffer C, and centrifuged at 72,000 *g* in a P28S swing rotor (Hitachi Koki) for 30 min at 4°C. The band at the 0-4% Ficoll interface was collected using a Piston Gradient Fractionator (BioComp Instruments, Inc). The vacuolar fraction was stored at -75°C. The yield of vacuole was estimated by measuring the enzymatic activity of the vacuolar enzyme Pho8, as previously described (Makino et al., 2021), or by calculating the band intensity of Pho8 detected by western blotting. Protease inhibitors and reducing agents were not used during the isolation of vacuoles.

### Fractionation of vacuolar lysate into soluble and pellet fractions

The vacuolar lysate was mixed with an equal volume of 2x buffer D and centrifugated at 135,000 *g* in a S55A2 angle rotor (Hitachi Koki) at 4°C for 30 min to obtain supernatant (*S*) fraction and pellet (*P*) fraction. *P* fraction was resuspended in buffer D containing either 2% TritonX-100 (Tx) (SIGMA), 0.5% DDM (SIGMA), 0.1 M Na_2_CO3, 2.5 M urea, or 1 M KCl and incubated for 30 min at 4 °C with shaking at 1,400 rpm. Then, the samples were centrifuged again at 135,000 *g* for 30 min at 4 °C to obtain a second supernatant (*S_2_*) fraction and pellet (*P_2_*) fraction.

### Immunoprecipitation and elution of processed forms of Atg15

Vacuolar pellet fractions obtained from 5 L yeast culture were suspended in 5 ml of buffer D containing 0.5% DDM by vortex and sonication, and then incubated with 60 μl (slurry 50% v/v) of α-FLAG antibody conjugated agarose beads (Millipore) for 1 h at 4°C. The beads were washed three times with buffer E and then mixed with 60 μg of 3xFLAG peptide (SIGMA) in 60 μl of buffer E for 1 h with shaking at 1400 rpm at 4°C. The samples were transferred to a Biospin column (BIORAD) and eluted by centrifugation. The volume of the eluate was approximately 60 μl from 5 L culture.

### In vitro lipase activity assay

To measure the lipase activity of vacuolar lysates, 125 ng of NBD-PE was added to each vacuolar lysate fraction in total volume of 20 μl of buffer B’ and incubated at 30°C for the indicated time periods. The amount of vacuolar lysate was normalized using either the amount of Pho8, as determined by the band intensity of Pho8 detected by western blotting, or the enzymatic activity of Pho8. Lipase assays of _prc_Atg15 eluates were performed by incubating samples with 125 ng of each fluorescent lipid (NBD-PE, NBD-PS, NBD-PC, NBD-PG, NBD-LPC, TF-PI, TF-TG, or BODIPY-CE) in a total volume of 20 μl buffer E. Total lipids of the reaction mixture were extracted according to the BUME method (Lofgren et al., 2012). The lipids were dissolved in chloroform/methanol (2:1, v/v), and applied to a TLC plate (HPTLC SILICA GEL 60 F254 50 GLASS PL 1.05642.0001, Millipore), which was developed with a solution containing chloroform/methanol/water at a ratio of 65:35:8 for NBD-PE NBD-PS, NBD-PC, NBD-PG, NBD-LPC, TF-PI or hexane/diethyl ether/acetic acid at a ratio of 50/50/1 for TF-TG and BODIPY-CE as described previously (Ishibashi et al., 2019). Fluorescence on the TLC plate was detected using a FUSION-FX7 imaging system (Vilber-Lourmat). The intensity of each fluorescent band was quantified using FUSION-FX7 analysis software.

### Isolation of ABs

ABs were isolated as described previously (Kawamata et al., 2022). In brief, *atg15*Δ cells (SMY345) were grown in 2.5 L YPD cultures to a density of OD_600_ = 1.0 and then treated with 0.2 μM rapamycin for 6 h. Vacuoles were isolated as described above, and then passed through a 0.8 μm PC membrane filter (ATTP01300, Millipore) to liberate ABs. Filtrates were subjected to 0 - 30% OPTIPREP density gradient centrifugation at 72,000 *g* for 90 min using a P40ST swing rotor and AB-enriched fractions (approximately 3 -4 ml) were collected.

### Evaluation of AB disruption in vitro

20 μl of AB fraction was mixed with 10 or 30 μl of _prc_Atg15 eluates or 0.2% Tx in total 50 μl of buffer F and incubated at 30°C for 3 h. Samples were then treated with 160 μg/ml proteinase K on ice for 30 min, and the reaction was terminated by adding PMSF. Samples were precipitated using TCA and analyzed by western blotting.

### Western blotting

Immunoblot analyses of Ape1 were performed as previously described (Urban et al., 2007). To analyze the processing of Atg15, the samples were precipitated with 10% of TCA and 1.8 μg/ml tRNA and resuspended in the sample buffer (50 mM Tris-HCl pH 7.5, 70 mM SDS, 8% Glycerol, 20 mM DTT, BPB), followed by incubation at 65°C for 15 min. For deglycosylation of Atg15, the samples were treated with Endo H (NEB) and GlycoBuffer (NEB) at 30°C for 2 h. Samples were then separated by SDS-PAGE and subjected to western blotting. α-FLAG (Sigma–Aldrich, F3165, 1:1000), α-Pho8 (Abcam, ab113688, 1:1000), α-Ape1 (1:5000) (Hamasaki et al., 2003), α-Prb1 (laboratory stock, 1:5000) and α-GFP (Roche, 11814460001, 1:1000), were used as primary antibodies. Femtoglow HRP Substrate (Michigan Diagnostics, 21008) was used to initiate chemiluminescence and blots were visualized using a FUSION-FX7 imaging system.

### Microscopic analysis

Fluorescence microscopy was performed using an inverted fluorescence microscope (Olympus, IX81) equipped with an electron-multiplying CCD camera (Hamamatsu Photonics, ImagEM C9100-13) and 150x objective lens (UAPON 150x OTIRF, NA/1.45; Olympus). mNeonGreen and mCherry were excited using 488 nm and 561 nm lasers, respectively, both with a maximum output of 50 mW (Coherent). Emitted fluorescence was filtered using a Di01-R488/561-25 dichroic mirror (Semrock) and an Em01-R488/568-25 bandpass filter (Semrock), and separated into two channels using a U-SIP splitter (Olympus) equipped with a DM565HQ dichroic mirror (Olympus). Fluorescence was further filtered using an FF02-525/50-25 bandpass filter (Semrock) for the mNeonGreen channel and an FF01-624/40-25 bandpass filter (Semrock) for the mCherry channel.

For super-resolution microscopy, a confocal microscope (IXplore SpinSR10) equipped with an sCMOS camera (ORCA-Flash4.0, Hamamatsu), a 100x objective lens (UPLAPO 100xOHR, NA/1.50; Olympus) was employed. mNeonGreen and mCherry were excited using 488 nm and 561 nm lasers, respectively. mNeonGreen was excited using a 488 nm laser, and fluorescence was passed through a mirror unit U-FGFP equipped with a DM490GFP dichroic mirror (Semrock), a BP460-480GFP excitation filter, and a BA495-540GFP absorption filter. mCherry was excited using a 561 nm laser and fluorescence passed through a U-FMCHE mirror unit (Olympus) equipped with a DM595 dichroic mirror, a BP565-585 excitation filter and a BA600-690 absorption filter. Images were acquired using super-resolution mode.

Images were acquired using MetaMorph (Molecular Devices) or CellSens (Olympus, IX81) software and processed using ImageJ software (Fiji Ver. 1.53f51) (Schindelin et al., 2012).

## Acknowledgements

We are grateful to members of the Ohsumi laboratory, Dr. Nobuo Noda, Dr. Yohei Ishibashi, and Dr. Masato Umeda for discussions and critical comments, and the Biomaterials Analysis Division, Open Facility Center, Tokyo Institute of Technology for DNA sequencing. This work was supported in part by Grants-in-Aid for Scientific Research 16H06375 and 19H05708 (to Y.O.) and 18H02399 (to T.K.) from the Ministry of Education, Culture, Sports, Science and Technology of Japan.

## Author contributions

Y.K., T.K. and Y.O. designed experiments and wrote the manuscript; Y.K., T.K. A.M. and M.S. performed the experiments.

The authors declare no conflicts of interest.

## Expanded view figure legends

**Figure EV1. Supplemental data of Figure 1**

(A) Purification of vacuole from WT cells. The whole-cell lysate (*T*) and vacuolar fraction (*V*) samples were subjected to western blot analysis using antibodies against Pho8 (vacuole membrane), Ape1 (autophagy cargo), Pep4, Prb1 (vacuole lumen), Pgk1, Hsp90 (cytosolic proteins), Dpm1 (ER membrane), Cox2 (mitochondrial matrix), Por1 (outer mitochondrial membrane), Van1 (Golgi), and Gsp1 (nucleus). Samples equivalent to 1/3000 of total lysate and 1/160 of vacuolar fraction were loaded.

(B) Western blotting of Pho8 in vacuolar lysates from WT, *atg15*Δ, and *atg15*Δ cells expressing Atg15 from either single or multi-copy (OE) plasmids. The band intensity ratio of Pho8 for each strain is shown under the panel.

(C) Western blotting of Pho8 in vacuolar lysates from *atg15*Δ cells expressing Atg15 or Atg15^S332A^ from multi-copy plasmids. The experimental procedure was the same as in (B).

(D) Western blotting of Pho8 in vacuolar lysates from WT, *prb1*Δ, *pep4*Δ, and *pep4*Δ *prb1*Δ cells. The experimental procedure was the same as in (B).

(E) Lipase activities of vacuolar lysates from WT, *pep4*Δ*prb1*Δ, and *pep4*Δ*prb1*Δ cells expressing Atg15 from a multi-copy plasmid. Each vacuolar lysate was added at a volume ratio of 1:5:25. The amount of the lysates used for the assay was shown in (F).

(F) Western blot of Pho8 in vacuolar lysates from WT, *atg15*Δ, and *pep4*Δ*prb1*Δ cells expressing Atg15 from multi-copy plasmids. The experimental procedure was the same as in (B).

**Figure EV2. Supplemental data of Figure 2**

(A) Western blotting of vacuolar fractions from WT, *atg15*Δ, and *atg15*Δ cells expressing C-terminally FLAG-tagged Atg15 (Atg15-FLAG) from either single or multi-copy plasmids. The ratio of the band intensities detected by α-FLAG (red squares) and α-Pho8 antibodies for each strain are shown in the right panels. *, nonspecific band.

(B) Lipase activity of vacuolar lysates from the strains in (A).

(C) Western blotting of vacuolar fractions from *atg15*Δ and *pep4*Δ*prb1*Δ cells each expressing Atg15-FLAG from a multi-copy plasmid. The experimental procedure was the same as in Fig EV1B.

(D) Maturation of prApe1 in mNeonGreen-tagged Atg15 expressing cells. Cells were grown in SD/CA medium and treated with rapamycin for 2 h. prApe1 and mApe1 were detected with α-Ape1 antibodies. Full length of C-terminally mNeonGreen-tagged Atg15 and CPY^N50^-Atg15^ΔN35^ were detected with α-mNeonGreen antibodies.

(E) Fluorescence microscopy image of mCherry-Atg8 in WT cells. Cells were grown in SD medium and then treated with rapamycin for 3 h. Scale bar, 1 μm. Dashed lines, vacuole boundaries.

(F) Western blotting of vacuolar fractions from *atg15*Δ and *pep4*Δ*prb1*Δ cells each expressing CPY^N50^-Atg15^ΔN35^. The experimental procedures were the same as in Fig. EV1B.

**Figure EV3. Supplemental data of Figure 4**

(A, B) Western blotting of the maturation of prApe1 in cells expressing N-terminally (A) or C-terminally (B) truncated Atg15 mutants. *atg15*Δ cells expressing WT Atg15 or fusions of truncated Atg15 with CPY^N50^ under the control of *ATG15* promoter from a multicopy-plasmid were cultured in the same conditions as in Fig 2D. A series of N- and C-terminally truncated forms of ATG15 were constructed by PCR from pRS426-CPY(1-50)- ATG15(Δ1-35)-mNeonGreen (YPL073) (see methods).

(C) Vacuolar pellet fraction (Fig 4C lane3) was treated with 2% Tx or 0.5% DDM (*T_2_*) and incubated at 4 °C for 30 min. Then samples were centrifuged at 135,000 *g* for 30 min, followed by separation into pellet (*P_2_*) and supernatant (*S_2_*) fractions before being subjected to western blot analysis with α-FLAG antibodies. *, nonspecific band.

(D) Vacuolar pellet fraction (Fig 4C lane3) was treated with 0.5% DDM or 2% Tx. Samples were then subjected to western blotting with α-FLAG antibodies. *, nonspecific band.

(E) Lipase activities of vacuolar pellet fraction treated with detergents. NBD-PE was incubated with each fraction in (D). Final DDM and Tx concentrations in the reaction mixture were 0.05% and 0.2%, respectively.

**Figure EV4. Supplemental data of Figure 5**

(A and B) Lipase activity of vacuolar lysates from *atg15*Δ cells expressing Atg15^iFLAG^ and Atg15^iFLAG,S332A^. The lysates were incubated with NBD-PE, NBD-PS (18:1-12:0), NBD-PC (18:1-12:0) and NBD-PG (18:1-12:0), each labeled at *sn-2* (A), and NBD-LPC (12:0) (B) for 1 h.

(C) Lipase activity of vacuolar lysates toward TF-PI (18:1-6:0) labeled at *sn-2*. The incubation time was 1 h.

(D) Lipase activity of vacuolar lysates toward TF-TG and BODIPY-CE. The incubation time was 2 h.

## Expanded view tables

**Expanded view table 1.** Yeast strains used in this study

| No. | Strain | Genotype | Source | Figures |
| --- | --- | --- | --- | --- |
| 1 | X2180-1B | *MATa SUC2 mal mel gal2 CUP1* | Yeast Genetic Stock Center | Fig EV1B, D-F, 2A, B, D |
| 2 | MMY209 | X2180-1B; *atg15Δ::kanMX6* | this study | Fig EV1B, 2A, B |
| 3 | KYY070 | X2180-1B; *atg15Δ::kanMX6, ura3Δ::hphNT1*<br>pRS426 | this study | Fig EV2D, 3A, B |
| 4 | KYY073 | X2180-1B; *atg15Δ::kanMX6, ura3Δ::hphNT1*<br>pRS426-*ATG15-mNeonGreen* | this study | Fig EV2D |
| 5 | SMY96 | X2180-1B; *ura3Δ::hphNT1, atg15Δ::kanMX6*<br>pRS426-*ATG15* | this study | Fig EV1B and C |
| 6 | SMY97 | X2180-1B; *ura3Δ::hphNT1, atg15Δ::kanMX6*<br>pRS426-*ATG15(S332A)* | this study | Fig EV1C |
| 7 | KYY127 | X2180-1B; *atg15Δ::kanMX6, ura3Δ::hphNT1*<br>pRS426-*CPY(1-50)-ATG15(Δ1-35)-mNeonGreen* | this study | Fig EV2D |
| 8 | KYY163 | X2180-1B; *atg15Δ::kanMX6, ura3Δ::hphNT1*<br>pRS426-*CPY(1-50)-ATG15(Δ1-55)* | this study | Fig EV3A |
| 9 | KYY164 | X2180-1B; *atg15Δ::kanMX6, ura3Δ::hphNT1*<br>pRS426-*CPY(1-50)-ATG15(Δ1-56)* | this study | Fig EV3A |
| 10 | KYY165 | X2180-1B; *atg15Δ::kanMX6, ura3Δ::hphNT1*<br>pRS426-*CPY(1-50)-ATG15(Δ1-57)* | this study | Fig EV3A |
| 11 | KYY166 | X2180-1B; *atg15Δ::kanMX6, ura3Δ::hphNT1*<br>pRS426-*CPY(1-50)-ATG15(Δ1-58)* | this study | Fig EV3A |
| 12 | KYY168 | X2180-1B; *atg15Δ::kanMX6, ura3Δ::hphNT1*<br>pRS426-*CPY(1-50)-ATG15(36-479)* | this study | Fig EV3B |
| 13 | KYY169 | X2180-1B; *atg15Δ::kanMX6, ura3Δ::hphNT1*<br>pRS426-*CPY(1-50)-ATG15(36-476)* | this study | Fig EV3B |
| 14 | KYY170 | X2180-1B; *atg15Δ::kanMX6, ura3Δ::hphNT1*<br>pRS426-*CPY(1-50)-ATG15(36-470)* | this study | Fig EV3B |
| 15 | KYY171 | X2180-1B; *atg15Δ::kanMX6, ura3Δ::hphNT1*<br>pRS426-*CPY(1-50)-ATG15(36-469)* | this study | Fig EV3B |
| 16 | KYY172 | X2180-1B; *atg15Δ::kanMX6, ura3Δ::hphNT1*<br>pRS426-CPY(1-50)-ATG15(36-468) | this study | Fig EV3B |
| 17 | KYY173 | X2180-1B; *atg15Δ::kanMX6, ura3Δ::hphNT1*<br>pRS426-CPY(1-50)-ATG15(36-466) | this study | Fig EV3B |
| 18 | KYY174 | X2180-1B; *atg15Δ::kanMX6, ura3Δ::hphNT1*<br>pRS426-CPY(1-50)-ATG15(36-465) | this study | Fig EV3B |
| 19 | KYY185 | X2180-1B; *pep4Δ::kanMX6, prb1Δ::hphNT1, leu2::ATG8pro-mCherry-ATG8(G116)-zeoNT3, ura3Δ::natNT2*<br>pRS426-CPY(1-50)-ATG15(Δ1-35) | this study | Fig EV2F |
| 20 | KYY187 | X2180-1B; *atg15Δ::kanMX6, ura3Δ::hphNT1*<br>pRS426-CPY(1-50)-ATG15(Δ1-59) | this study | Fig EV3A |
| 21 | KYY188 | X2180-1B; *atg15Δ::kanMX6, ura3Δ::hphNT1*<br>pRS426-CPY(1-50)-ATG15(Δ1-60) | this study | Fig EV3A |
| 22 | KYY193 | X2180-1B; *atg15Δ::kanMX6, ura3Δ::hphNT1*<br>pRS426-CPY(1-50)-ATG15(36-467) | this study | Fig EV3B |
| 23 | KYY194 | X2180-1B; *atg15Δ::kanMX6, ura3Δ::hphNT1*<br>pRS426-CPY(1-50)-ATG15(Δ1-35) | this study | Fig EV3A and B |
| 24 | KYY201 | X2180-1B; *atg15Δ::kanMX6, ura3Δ::hphNT1, leu2::ATG8pro-mCherry-ATG8(G116)-zeoNT3*<br>pRS426-CPY(1-50)-ATG15(Δ1-35) | this study | Fig EV2F |
| 25 | SMY96 | X2180-1B; *ura3Δ::hphNT1, atg15Δ::kanMX6*<br>pRS426-ATG15 | this study | Fig EV2D |
| 26 | SMY125 | X2180-1B; *atg15Δ::kanMX6, ura3Δ::hphNT1*<br>pRS426-ATG15-3xFLAG | this study | Fig EV2A-C |
| 27 | SMY127 | X2180-1B; *ura3Δ::natNT2, prb1Δ::kanMX6, pep4Δ::hphNT1*<br>pRS426-ATG15 | this study | Fig EV1E and F |
| 28 | SMY260 | X2180-1B; *ura3Δ::hphNT1, atg15Δ::kanMX6*<br>pRS316-ATG15 | this study | Fig EV1B |
| 29 | SMY261 | X2180-1B; *atg15Δ::kanMX6, ura3Δ::hphNT1*<br>pRS316-ATG15-3xFLAG | this study | Fig EV2A and B |
| 30 | SMY311 | X2180-1B; *pep4Δ::kanMX6, prb1Δ::hphNT1* | this study | Fig EV1D-F |
| 31 | SMY476 | X2180-1B; *atg15Δ::kanMX6, ura3Δ::natNT2, ape1Δ::zeoNT3*<br>pRS426-ATG15-3xFLAG | this study | Fig EV2C |
| 32 | SMY485 | X2180-1B; *pep4Δ::hphNT1*, *prb1Δ::kanMX6*,<br>*ura3Δ::natNT2*, *ape1Δ::zeoNT3*<br>pRS426-ATG15-3xFLAG | this study | Fig EV2C |
| 33 | SMY491 | X2180-1B; *atg15Δ::kanMX6*, *ura3Δ::hphNT1*,<br>*ape1Δ::natNT2*<br>pRS426-ATG15(168-3xFLAG-169) | this study | Fig EV3C-E, EV4 |
| 34 | SMY492 | X2180-1B; *atg15Δ::kanMX6*, *ura3Δ::hphNT1*,<br>*ape1Δ::natNT2*<br>pRS426-ATG15(S332A, 168-3xFLAG-169) | this study | Fig EV4 |
| 35 | SMY649 | X2180-1B; *ura3Δ::hphNT1*, *leu2::ATG8pro-mCherry-ATG8(G116)-zeoNT3*<br>pRS426 | this study | Fig EV2E |
| 36 | ScYO517 | X2180-1B; *prb1Δ::hphNT1* | this study | Fig EV1D |
| 37 | MMY110 | X2180-1B; *pep4Δ::hphNT1* | this study | Fig EV1D |

**Expanded view table 2.** Plasmids used in this study

| No. | Plasmids | Details | source |
| --- | --- | --- | --- |
| 1 | YPL111 | pRS426-CPY(1-50)-ATG15(36-469) | this study |
| 2 | YPL112 | pRS426-CPY(1-50)-ATG15(36-479) | this study |
| 3 | YPL114 | pRS426-CPY(1-50)-ATG15(36-466) | this study |
| 4 | YPL116 | pRS426-CPY(1-50)-ATG15(36-476) | this study |
| 5 | YPL117 | pRS426-CPY(1-50)-ATG15(36-470) | this study |
| 6 | YPL119 | pRS426-CPY(1-50)-ATG15(36-468) | this study |
| 7 | YPL120 | pRS426-CPY(1-50)-ATG15(36-465) | this study |
| 8 | YPL122 | pRS426-CPY(1-50)-ATG15(Δ1-55) | this study |
| 9 | YPL124 | pRS426-CPY(1-50)-ATG15(Δ1-56) | this study |
| 10 | YPL126 | pRS426-CPY(1-50)-ATG15(Δ1-57) | this study |
| 11 | YPL127 | pRS426-CPY(1-50)-ATG15(Δ1-58) | this study |
| 12 | YPL131 | pRS426-CPY(1-50)-ATG15(Δ1-59) | this study |
| 13 | YPL133 | pRS426-CPY(1-50)-ATG15(Δ1-60) | this study |
| 14 | YPL142 | pRS426-CPY(1-50)-ATG15(36-467) | this study |
| 15 | TPL004 | pRS316-ATG15-3xFLAG | this study |

**Expanded view table 3.**
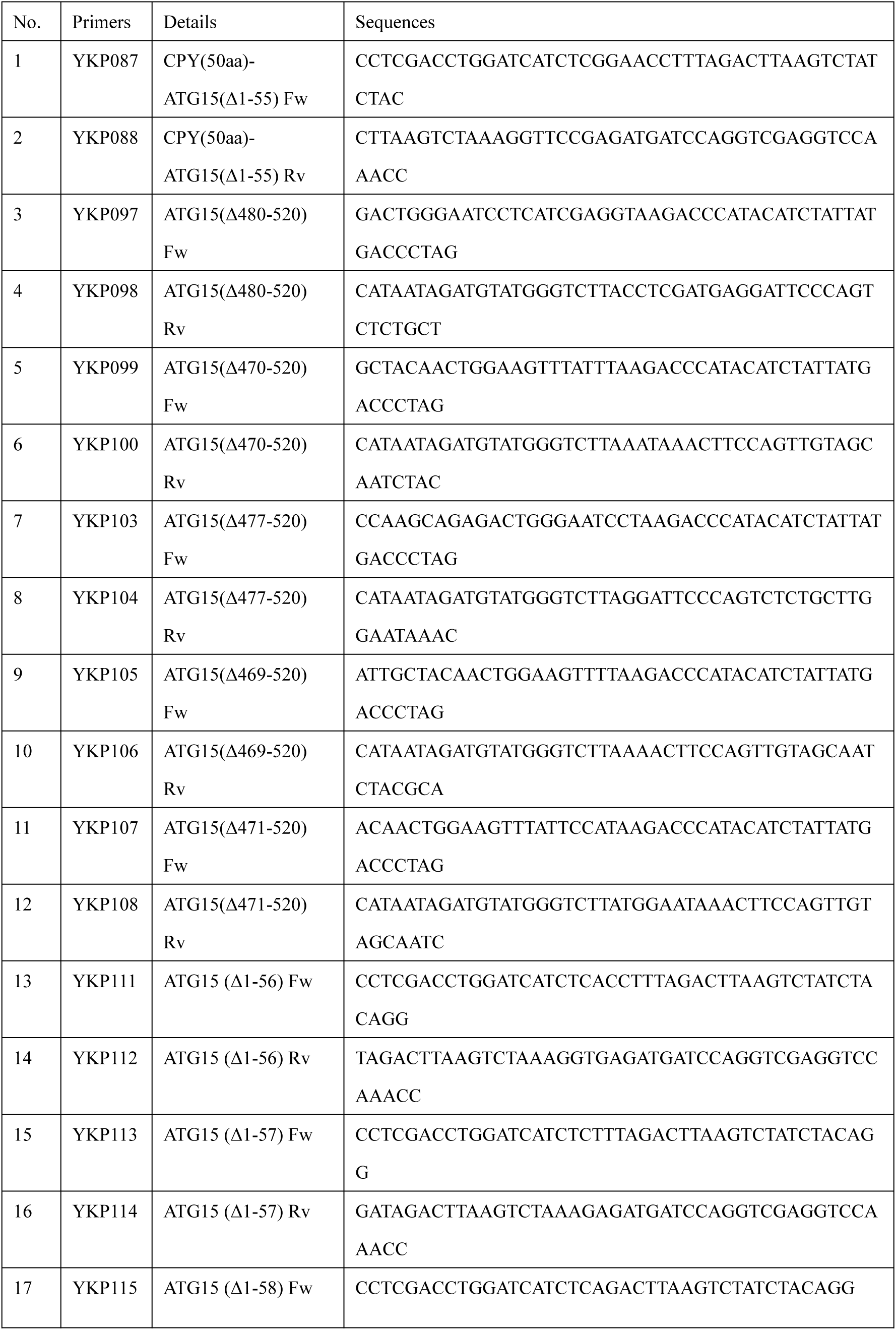
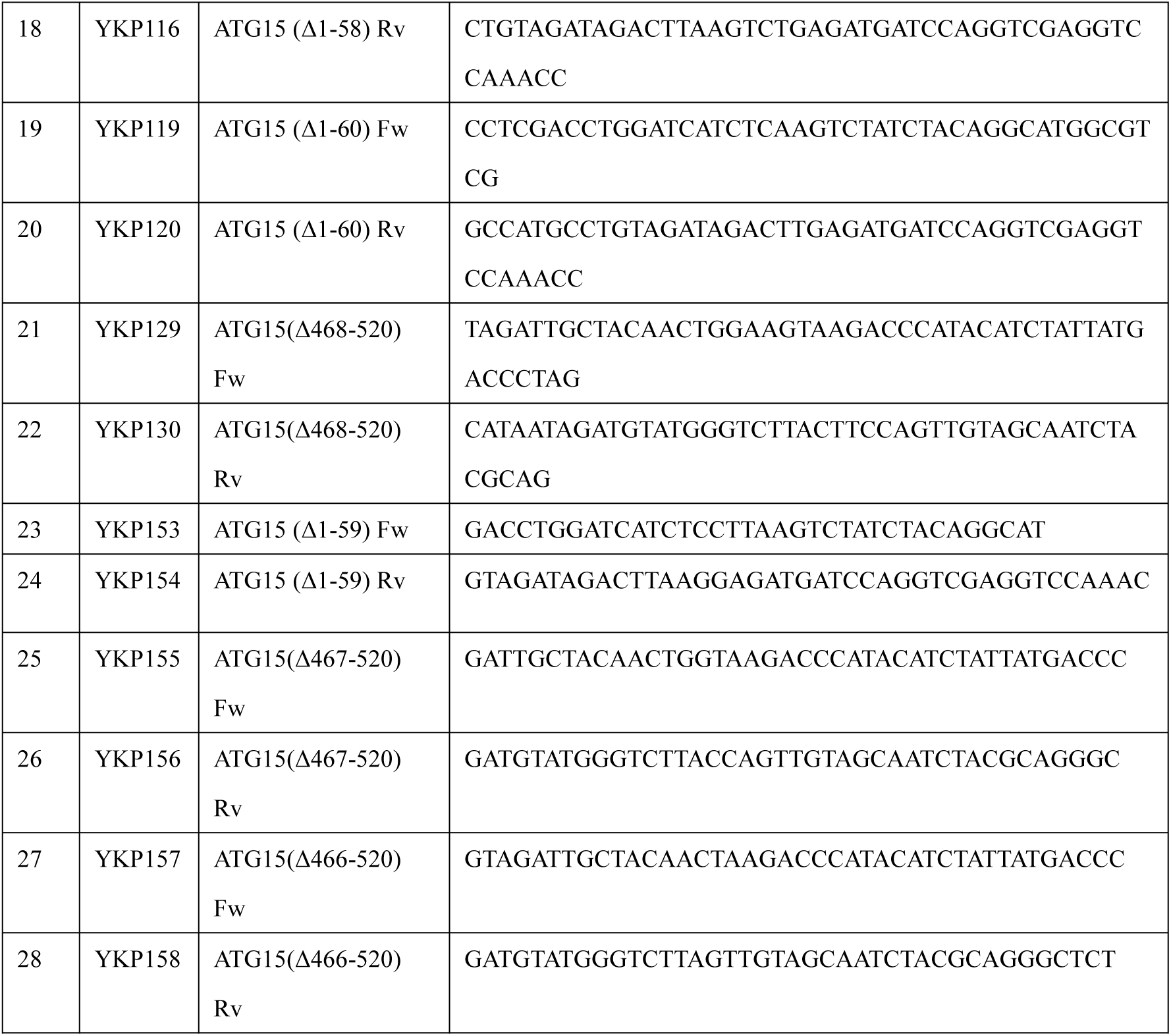
Primers used in this study

