## supplemental Figures for "The mechanism of Atg15-mediated membrane disruption in autophagy"

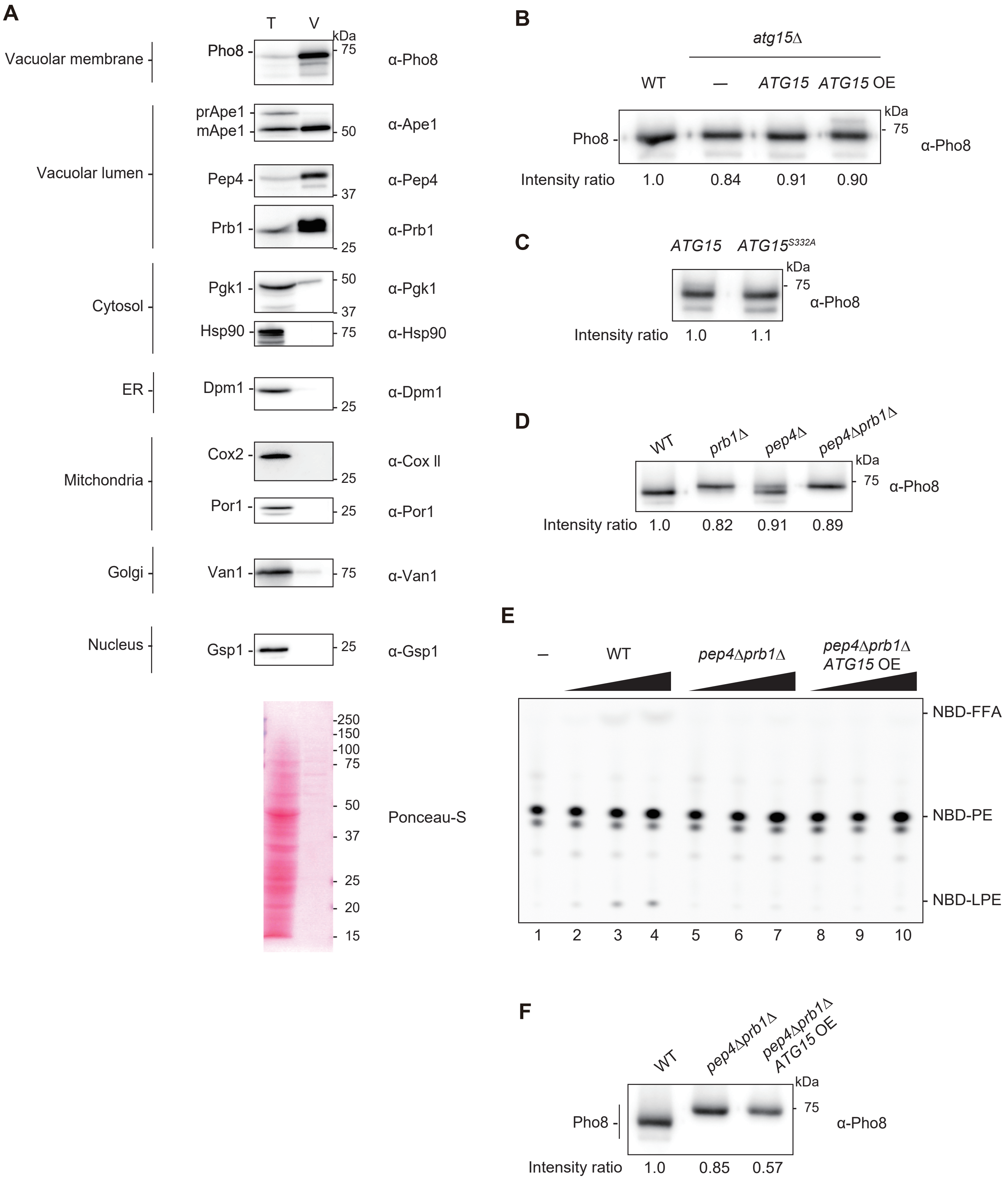


**Figure EV1. Supplemental data of Figure 1**

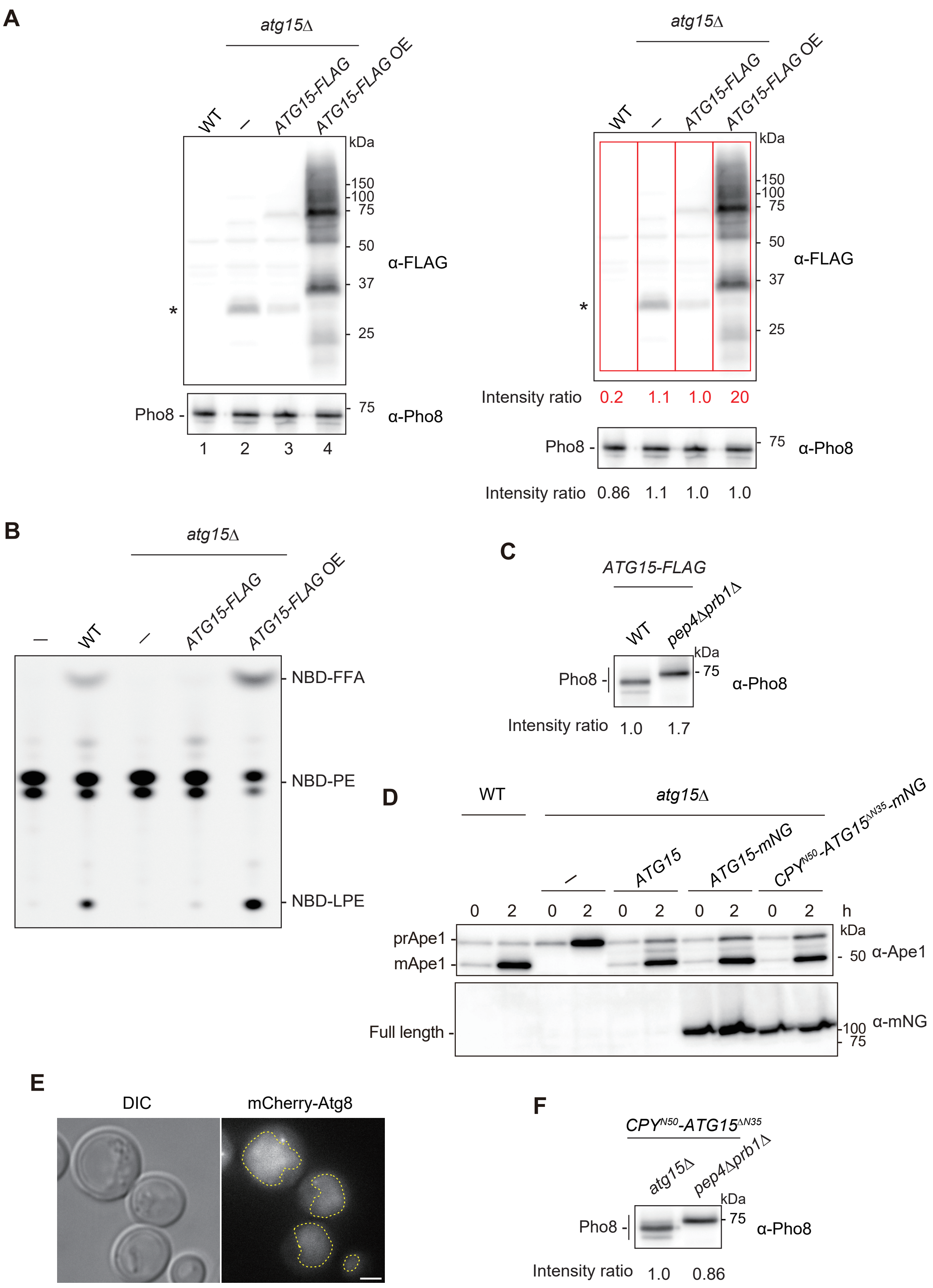


**Figure EV2. Supplemental data of Figure 2**

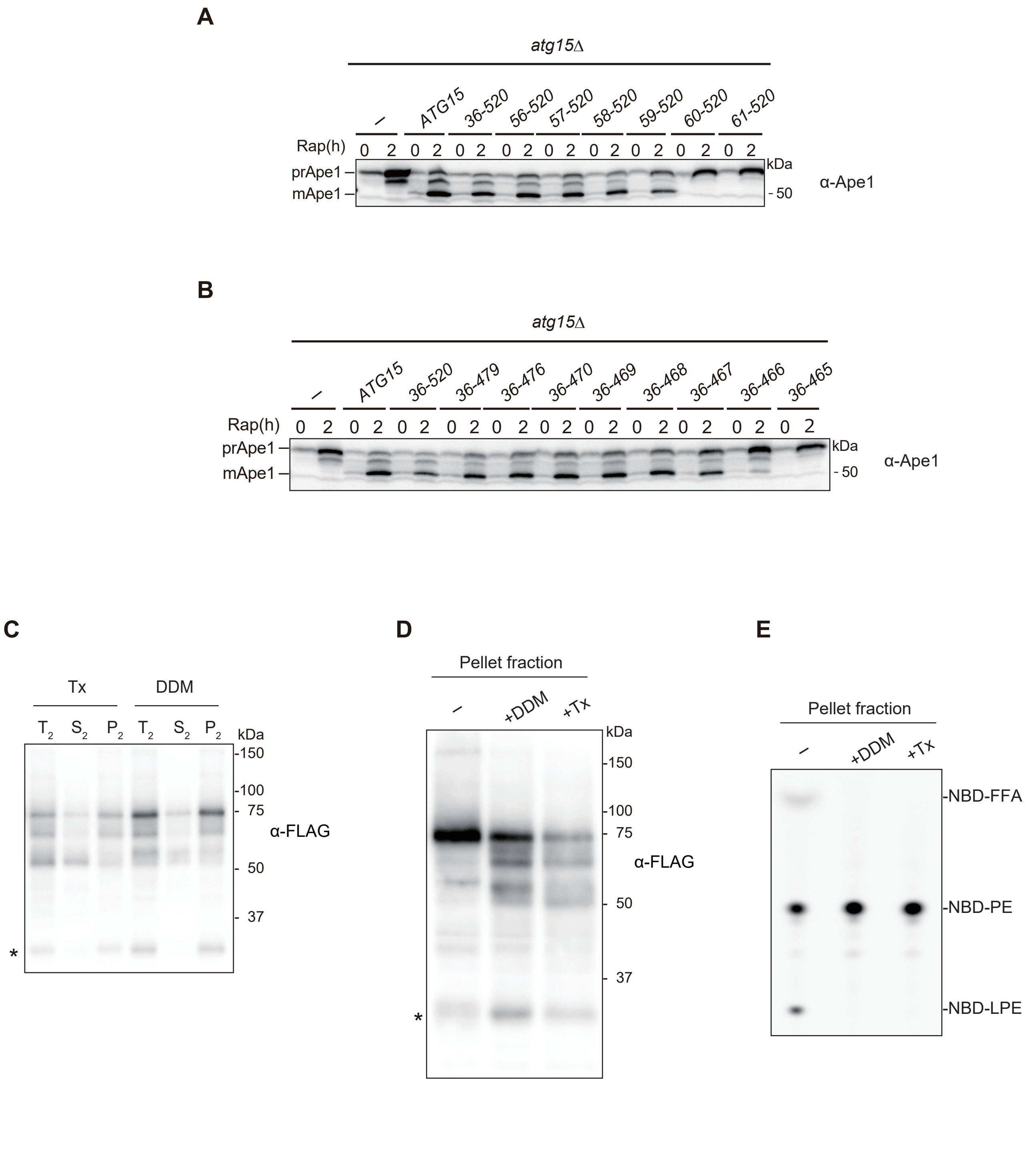


**Figure EV3. Supplemental data of Figure 4**

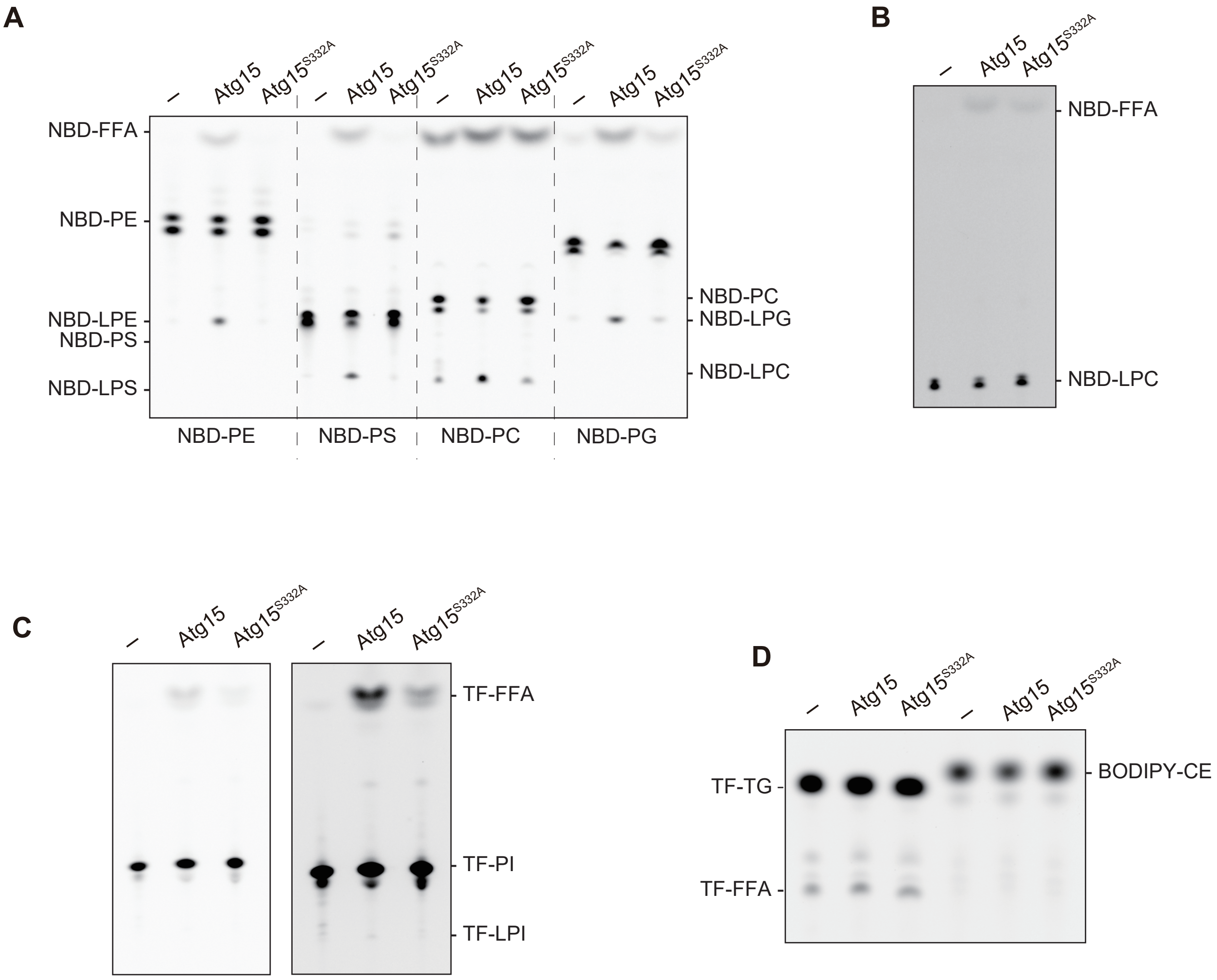


**Figure EV4. Supplemental data of Figure 5**
